# Learning from Hi-C reveals nuclear dynamical reorganization in budding yeast across the cell cycle

**DOI:** 10.64898/2026.11.07.764600

**Authors:** Antonio B. Oliveira, Vinícius G. Contessoto, Julien Mozziconacci, José N. Onuchic, Vittore F. Scolari

## Abstract

Hi-C contact maps provide population-averaged measurements of chromosomal contacts. Converting these measurements, however, into three-dimensional structures and associated dynamics remains an underdetermined problem. Existing approaches broadly fall into two classes. Data-driven reconstruction methods that infer either a consensus structure or an ensemble of conformations, but do not inform on chromosome dynamics. Physics-based models, which generate dynamical trajectories, but require prior assumptions about nuclear geometry, chromatin mechanics, tethering, or molecular interactions. Here, we bridge these approaches using the Full-Inversion Chromatin model, a data-driven whole-genome polymer framework that learns an effective force field directly from Hi-C contact probabilities; integrated with Serpentine, a tool to quantify contact-map discrepancies while accounting for differences in sequencing coverage and sparsity. This generates structural ensembles of chromosomes and effective chromatin dynamics. We apply this method at 5-kb resolution to *Saccharomyces cerevisiae* in G1, mid-S, S/G2, G2/M, anaphase, and quiescence. Without explicitly imposing the spindle pole body, centromere or telomere tethering, predefined chromatin states, or nuclear-envelope interactions, the inferred ensembles recover these major features of yeast nuclear organization. Importantly, although dynamical information is not used during model training, the resulting polymer trajectories reproduce the subdiffusive scaling and chromosomal variations of independently measured chromatin motion. The model further predicts heterogeneous mobility along chromosomes and a distinctive telomere-dependent dynamical organization. Thus, population-averaged Hi-C contacts contain sufficient information to infer not only three-dimensional genome organization, but also effective, locus-dependent chromatin dynamics.

## 1 Introduction

Translating Hi-C contact maps into three-dimensional genome organization is intrinsically underdetermined. Similar population-averaged contact frequencies can arise from different structures, conformational fluctuations, and nuclear constraints. Computational approaches to this inverse problem can broadly be divided into data-driven and physics-based strategies.

Data-driven methods seek conformations that reproduce the experimental contact matrix, either as a single consensus structure or as a heterogeneous ensemble. Consensus reconstructions provide a compact representation of genome folding [1, 2], whereas population-based approaches explicitly account for structural heterogeneity and the probabilistic nature of Hi-C contacts [3–5]. Data-driven inference has also been combined with polymer physics to infer effective interaction models that generate structural ensembles [6–12]. Such models can predict chromatin dynamics [13], but these predictions require assumptions about the underlying dynamical processes that are not uniquely determined by population level contact frequencies.

Physics-based polymer models generate explicit dynamical trajectories and allow mechanistic hypotheses to be tested. Their predictions, however, depend on prior assumptions about chromatin mechanics, nuclear geometry, confinement, or locus-specific interactions. A central challenge is therefore to infer a physically generative model from Hi-C data while minimizing external structural assumptions. This also raises a key question: does the inferred interaction landscape contain sufficient information to predict the chromatin dynamics that were not used for model inference?

Budding yeast provides a stringent system in which to address this challenge. Its nuclear architecture and chromatin mobility have been extensively quantified through imaging, Hi-C, and polymer modeling [14–17]. Its genome comprises 16 linear chromosomes totaling approximately 12.1 million base pairs [18–20]. Physics-based whole-nucleus models explain major features of its Rabl configuration through chromosome connectivity, centromere attachment to the spindle pole body, telomere association with the nuclear envelope, nucleolar exclusion, and confinement [21–23]. The Yeast Standard Model (YSM) implements these constraints in coarse-grained molecular dynamics (MD) simulations, providing a physical description of whole-genome architecture and chromatin dynamics [16, 17, 23, 24].

The YSM also serves as a testbed for molecular mechanisms. Extensions have addressed telomere anchoring and clustering [25], cohesin activity and chromosome replication [26, 27], and Rad51-mediated homology search [28], yielding predictions that guide and interpret experiments. YSM simulations have also provided insights into nucleolar structure, supported by fluorescence imaging [29]. These models prescribe structural constraints from prior biological knowledge. The YSM was developed and validated primarily in the G1 phase of the cell cycle of exponentially growing cells, drawing on extensive quantitative measurements of yeast chromosome organization. These include Hi-C contacts, three-dimensional locus positions, distances between pairs of loci, and chromatin dynamics quantified by mean-square displacements [14, 30–33]. Its application to genome organization throughout the full cell cycle therefore remains an open challenge.

Yeast offers a rare opportunity to address both limitations through a single question: how much of its three-dimensional genome organization and chromatin dynamics can be recovered across the full cell cycle from Hi-C contacts, without prescribing nuclear landmarks or using dynamical measurements for model inference?

Here we advance beyond this established framework and take a complementary approach to this problem. We use machine-learning to invert the mapping from a population of chromosomal structures to Hi-C contact maps [12], and integrate Hi-C data across the entire yeast cell cycle. The method infers a polymer Effective Force Field (EFF) compatible with Hi-C contact data, assuming only the size of individual chromosomes and their chromatin backbone connectivity. It thereby enables the generation of an ensemble of full three-dimensional chromosome conformations, together with their interchromosomal interactions, that is consistent with the observed Hi-C data. Using a set of Hi-C contact maps spanning the entire yeast cell cycle [34, 35], we apply this framework to model the temporal evolution of chromosome structure. The model generates quantitative, falsifiable predictions and identifies experimental conditions and measurements that can enable a more detailed characterisation of yeast chromatin conformation and its dynamics throughout the cell cycle.

## 2 Methods

### 2.1 Budding yeast whole-genome polymer model

The *Saccharomyces cerevisiae* genome was modeled as a system of 16 chromosomes, represented as coarse-grained polymers at a 5-kb resolution per bead. Because the *rDNA* region in chromosome XII is not resolved in Hi-C experiments, chromosome XII was split into two independent polymer chains by removing the *rDNA* segment from the simulation model. As a consequence, the complete budding yeast genome was represented by 17 polymer chains evolved simultaneously in the same simulation box.

The bonded terms of the model were described by the same classical homopolymer potential used in the Full-Inversion Chromatin (FI-Chrom) framework [12], which includes bond connectivity, local bending rigidity, an excluded-volume interactions. The explicit functional form of these terms is described in the SI. The non-bonded interactions were inferred directly from Hi-C data using the FI-Chrom, in which all pairwise contact restraints are encoded through an EFF learned from the experimental contact probabilities.

### 2.2 Full-Inversion Chromatin non-bonded interaction

The FI-Chrom model is a data-driven maximum-entropy framework in which the total potential energy is written as

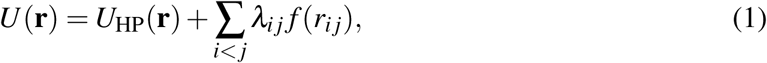

where *U*_HP_(r) is the homopolymer contribution, λ*_i j_* are the effective pairwise interaction parameters learned from Hi-C, and *f* (*r_i j_*) is a smooth contact function that maps spatial distance into contact probability. In this formulation, the structural ensemble is minimally biased beyond the information contained in the experimental contact map, as required by the maximum-entropy principle [36].

The contact function was represented by a smooth switching function,

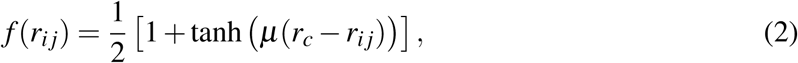

where *r_i j_* is the spatial distance between beads *i* and *j*, and µ and *r_c_* control the sharpness and effective contact cutoff, respectively.

### 2.3 FI-Chrom training pipeline

The FI-Chrom training protocol followed the same iterative simulation–optimization strategy described in the original framework [12], but here extended to the budding yeast whole genome. The optimization spans both intra-chromosomal and inter-chromosomal interactions, capturing the coupled organization of all chromosomes inside the same nucleus.

To improve training efficiency and stabilise the learning of the contact landscape, the Serpentine procedure was incorporated into the original pipeline [37]. Serpentine quantifies discrepancies between predicted and experimental contact maps by optimally binning data according to the sequencing coverage and returning the differential map

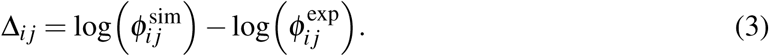

Serpentine was applied individually to intra-chromosomal regions and to each pair of chromosomes for inter-chromosomal regions, and the resulting maps were then combined. We took as a metric ∑*_i j_* |Δ*_i j_*| and optimized it in the iterative procedure. This facilitates the simultaneous inference of intra-and interchromosomal interactions, whose differing levels of sparsity are explicitly handled by the algorithm.

Briefly, experimental contact probabilities were first extracted from the Hi-C maps, and an initial force-field matrix λ^(0)^_*ij*_ was constructed. Simulations were then performed using the current force field to estimate the contact probabilities generated by the model, φ^sim^_*ij*_. The discrepancies from the experimental values φ^exp^_*ij*_, quantified by the metric defined above, were used to update the interaction parameters by the Adam optimizer [38]. This simulation → update → simulation cycle was repeated until the contact map error metric converged.

### 2.4 Generation of contact maps from molecular dynamics simulations

Contact maps were calculated from the ensemble of three-dimensional genome configurations generated by the FI-Chrom simulations. Molecular dynamics simulations were performed using Open-MiChroM [12], a chromatin simulation framework built on the OpenMM molecular dynamics engine [39]. For each pair of genomic loci *i* and *j*, the instantaneous probability of contact was defined using the smooth switching function:

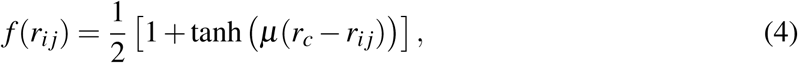

where *r_i j_* is the spatial distance between beads *i* and *j*, *r_c_* is the characteristic contact distance, and µ controls the sharpness of the transition between contacting and non-contacting configurations. We used µ = 2.0 and *r_c_* = 2.0, following the contact definition used in our previous chromatin simulations.

The simulated contact probability between loci *i* and *j* was obtained by averaging the contact function over the sampled conformational ensemble,

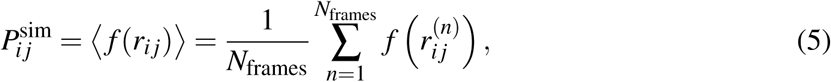

where 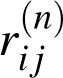 is the distance between loci *i* and *j* in simulation frame *n*, and *N*_frames_ is the total number of configurations included in the analysis. Thus, *P*^sim^ varies continuously between zero and one and represents the fraction of the conformational ensemble in which the two loci are spatially proximal, weighted by the smooth contact function.

For each cell-cycle stage, contacts were calculated for all pairs of loci across the complete simulated yeast genome, including both intra-chromosomal (*cis*) and inter-chromosomal (*trans*) pairs. The resulting whole-genome contact matrix therefore contains intra-chromosomal contacts along the diagonal blocks and inter-chromosomal contacts in the off-diagonal blocks. Contact maps were constructed at the same 5 kb resolution used in the polymer representation, allowing direct comparison with the corresponding experimental Hi-C maps.

### 2.5 Cell-cycle Hi-C datasets

To investigate cell-cycle-dependent genome organization, the model was trained on six independent budding yeast Hi-C datasets: G1, Mid-S, S/G2, G2/M, anaphase, and G0 (starvation) from a published dataset comprising the yeast full cell cycle [34, 35]. These are accessible at the NCBI GEO database, with accession numbers: GSM2417263 (G0), GSM2417264 (G1), GSM2417266 (Mid-S), GSM2417267 (S/G2), GSM2417268 (G2) and GSM2417271 (anaphase). For each stage, a separate FI-Chrom force field was inferred from the corresponding experimental contact map, allowing stage-specific changes in both intra-and inter-chromosomal interactions to be captured.

## 3 Results

### 3.1 Whole-genome modeling can recover high-fidelity chromosome features

Our machine learning approach (described in Methods 2.3) is based on the Full-Inversion Chromatin algorithm (FI-Chrom) and the Serpentine tool to quantify contact-map discrepancies while accounting for differences in sequencing coverage and sparsity [12, 37]. It was applied to the study of Yeast conformation in G1. The analysis was performed at a coarse-grained resolution of 5 kb, matching that of the contact maps. Each 5-kb bead represents approximately 27 nucleosomes. The training starts with a homopolymer potential that is modified by the inclusion of an energy function trained on experimental data using the maximum entropy principle [36, 40–43]. Constraints are derived as observable quantities from Hi-C maps used for optimization [7, 44, 45].

The trained model generates a molecular dynamics simulation of the chromatin conformation, that is used to generate three-dimensional snapshots similar to the example shown in Fig. 1A, with each chromosome colored according to the legend. These conformations exhibit chromosome territorialization: in an individual snapshot, each chromosome is confined to a limited region of the nucleus rather than spanning the full nuclear volume. Nevertheless, chromosomal territories belonging to different chromosomes are not exclusively defined domains in three dimensions, and intermix visually with each other. This observation is analogous to what has been reported in mammal cell lines, where interchromosomal intermingling happens principally in high gene density regions [46, 47].

**Figure 1:**
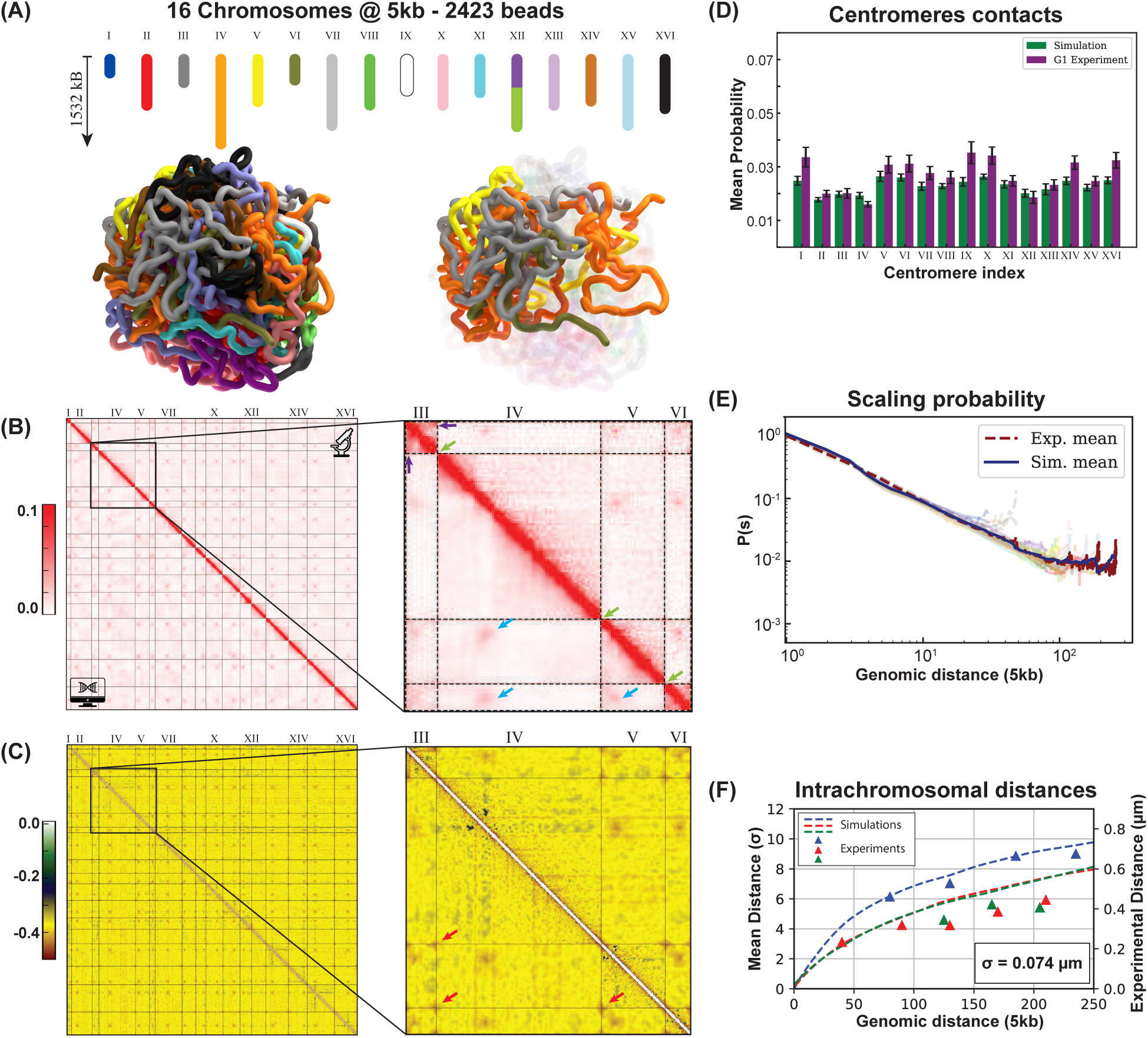
Whole-genome FI-Chrom model of budding yeast in G1. (A) Chromosome representation at 5-kb resolution, comprising 2423 beads, and a representative genome conformation with a zoom-in on chromosomes III–VI. Chromosomes are colored by index; chromosome XII is split at the unresolved rDNA region. (B) Experimental (upper triangle) and simulated (lower triangle) contact maps. Arrows indicate chromosome boundaries (green), centromere contacts (blue), and the interaction between mating-type loci on chromosome III (purple). (C) Inferred effective forcefield matrix, with negative and positive λ*_i j_* values indicating attractive and repulsive contributions, respectively. Red arrows mark telomeric interaction signatures. (D) Mean pericentromeric contact probabilities from Hi-C (green) and simulations (purple). (E) Mean intrachromosomal contact probability *P*(*s*) as a function of genomic separation. (F) Inter-locus distances from imaging [23] (triangles) and simulations (dashed lines). Colors distinguish pairs involving a locus 5 kb (blue), 854 kb (green), or 1185 kb (red) from the centromere.

Contact maps have been measured on simulated ensembles (see Methods). At the full genome level, the chromosome boundaries are identified (Fig. 1B, green arrows), as well as the clustering of the centromeres that is a signature of the SPB (Fig. 1B, blue arrows). The interaction between the centromere of one chromosome and any other centromeres was quantified, closely reproducing, in simulations, what is observed in contact maps (Fig. 1D). In contrast to the YSM, our approach is explicitly unbiased but does not allow testing mechanistic hypotheses. From a modeling perspective, the SPB is a 16-body interaction between all yeast point centromeres in the YSM, while only binary interactions are retained in our model. Results highlight how these two models are indistinguishable in Hi-C contact maps.

At the intra-chromosomal scale, the loop between the mating recombination sites on chromosome III, first described by Hi-C analysis [48], is identified (Fig. 1B, purple arrow). Further, the number of contacts generally decreases as a function of the genomic distance. The *P*(*s*) function quantifies the expected probability of contact between beads *i* and *i* + *s* in function of their genomic distance *s* based on the mean scaling of secondary diagonals in the contact map. This is reproduced accurately for the mean across all chromosomes (Fig. 1E). Taking advantage of our generated three dimensional conformations, we are now able to also quantify the scaling of the mean distance of two fluorescent operators on beads *i* and *i* + *s* averaged over a population ensemble, in function of their distance *s*:

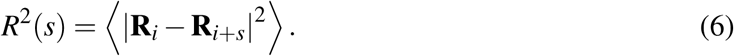

This features a sublinear scaling compatible with a generic polymer model. Comparing simulations to the published experimental data [23] allowed to fix the ratio between simulation spatial units σ and real spatial units, resulting in value of 74 nm/σ. Importantly, while contact maps can serve as a proxy for relative distances, they do not allow such a mapping without integrating imaging data. The model fitted parameters (λ*_i_*, *_j_*) are contained in the EFF, which quantify the level of attraction for each pairwise interaction (see Methods). EFFs for the G1 condition are plotted as a matrix in (Fig. 1C). In the YSM, nuclear confinement keeps chromosomes close enough to form contacts. Our model has no explicit nuclear boundary; instead, fitting the Hi-C contact map assigns generally negative values to λ*_i_*, *_j_*. These effective attractions thus capture the effect of confinement thas is learnt from the Hi-C data. In the phenomenological sense, confinement and general bead attraction can produce similar polymer organization [49]. The polymer *R*(*s*) and *P*(*s*) scalings are not affected, because the scaling of an equilibrated polymer mesh and a homopolymer in a good solvent are the same [50]. Beyond this general effect, we find Hi-C interaction signatures at telomeres in the EFF (red arrows) that are not typically considered in the YSM. These interactions, that emerge naturally in our model, are known to be dynamic in interphase, where they have been observed in microscopy [51–53], but play a crucial role in quiescence where they were previously observed in Hi-C contact maps [25, 34, 54–57].

Overall, our approach provides an unbiased, machine-learning-based generative framework for inferring three-dimensional chromosome conformations solely from Hi-C data acquired from yeast cells synchronized in G1 by elutriation [58]. This framework not only reproduces the main features of the contact probability maps, but also predicts three-dimensional conformations and mean interlocus distances.

### 3.2 Structural features across the cell cycle

The same procedure described in the previous section was applied to Hi-C libraries obtained from cells synchronized at different stages of the cell cycle, namely G1, mid-S, S/G2, G2/M, anaphase, and G0 (starvation), using the published dataset [34, 35]. Representative three-dimensional structures inferred from these datasets are shown in Fig. 2A. Because each Hi-C matrix gives rise to a distinct EFF, the resulting three-dimensional conformations can be used to characterize how yeast genome organization evolves across the full cell cycle.

**Figure 2:**
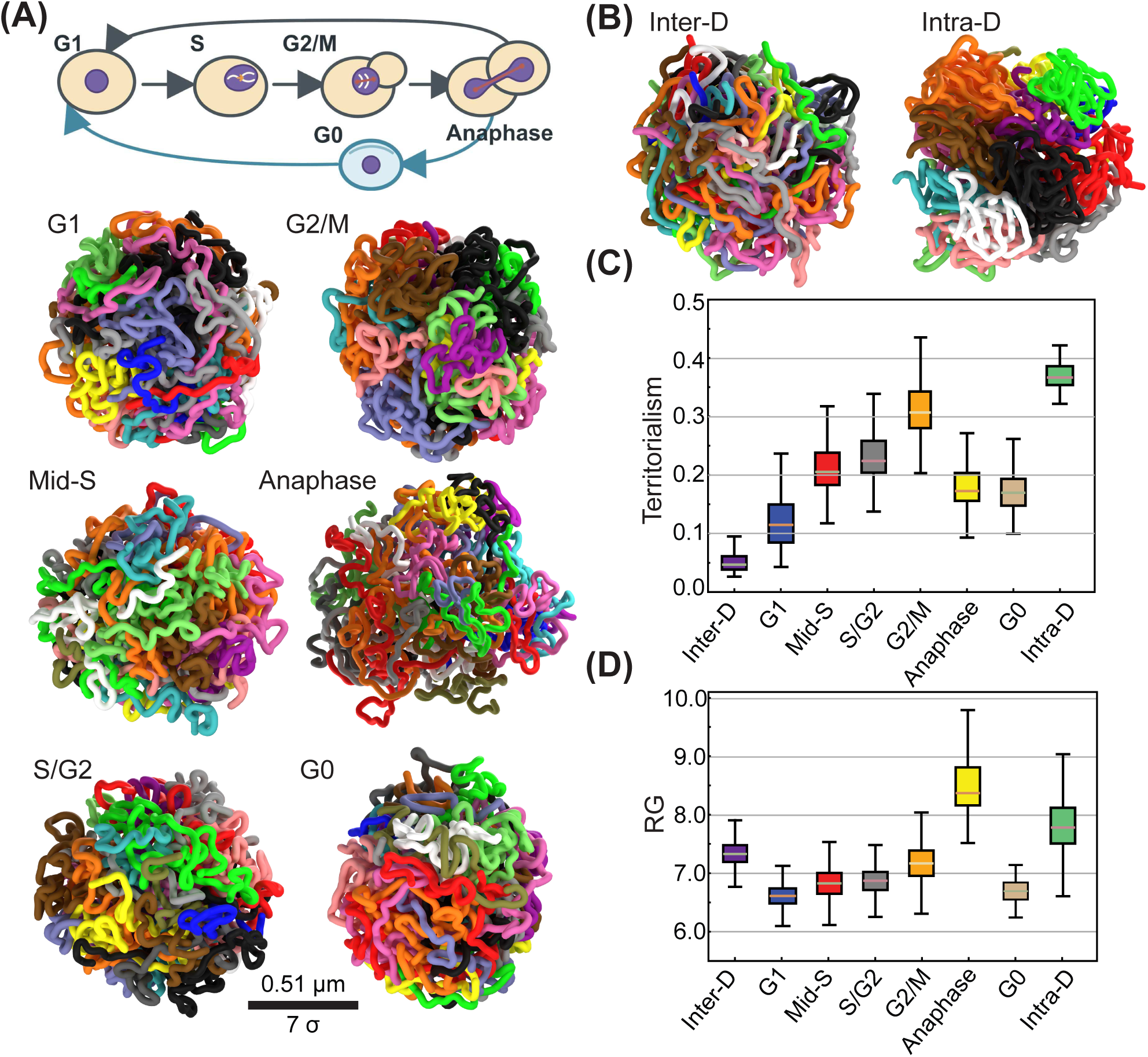
Structural characterization of the modeled genome across the cell cycle. (A) Schematic of the budding yeast cell cycle and representative genome conformations for G1, mid-S, S/G2, G2/M, anaphase, and G0. Chromosomes are colored as in Fig. 1A. (B) Representative conformations of the Intra-Dominant (Intra-D) and Inter-Dominant (Inter-D) reference models. (C) Chromosome territoriality quantified from intra-and interchromosomal distances across ensembles of 200,000 conformations. (D) Radius of gyration across the modeled conditions.

Visual inspection indicates that chromosomes in the G2/M condition are more spatially segregated, and therefore less intermixed, than in G1. This difference is consistent with the increased degree of chromosomal compaction due to the cohesin-mediated loops in metaphase [35, 59]. As a result, individual chromosomes become more clearly distinguishable. Although the metaphase plate is a hallmark of metazoan mitosis, the organization we infer in yeast is consistent with the analogous expectation that chromosomes adopt a more ordered and separated arrangement at metaphase to ensure faithful segregation. Next chromosome territoriality was quantified across all conditions and reported in Fig. 2C. The results indicate a progressive increase in cohesin-dependent territorial organization from G1 to G2/M, consistent with the gradual accumulation and action of cohesins throughout replication. In contrast, the anaphase and G0 conditions exhibit intermediate behavior. In budding yeast, G1 during exponential growth is a short-lived interphase state that follows chromosome segregation and precedes renewed DNA replication happening in S phase. This confirms observations that while cohesin is loaded in G1, loop extrusion and cohesion progresses during S phase [60].

The compaction and territoriality measured across cell-cycle stages is compared with two limiting reference models that were prescribed *ad hoc* to provide null references to the level of interchromosomal over intrachromosomal interactions. A inter-dominant (Inter-D) and a intra-dominat homopolymer (Intra-D). In both models, chromosomes are represented as chains of identical beads and therefore neglect most specific structural features of chromatin organization. The two models differ only in the relative strength of intra-chromosomal and inter-chromosomal bead–bead attractions (See Supplementary Fig. S6). In the Inter-Dominant model (Intra-D), all beads attracts each others just the same, whereas in the Intra-Dominant (Inter-D) model intra-chromosomal attractions are prescribed stronger than inter-chromosomal ones. Representative structures for these models are shown in Fig. 2B, and their corresponding EFFs are reported in Supplementary Fig. S1–S5.

In Fig. 2C we show that, with respect to territoriality, the G1 conformations are closest to the Inter-D regime, while the stronger compaction and individualization observed in G2/M shift the system toward the Intra-D regime. This is consitent with progressive cohesin loading and looping in S phase functional for the resolution of entanglements and chromosome segregation.

Focusing on the anaphase condition, visual inspection suggests that the genome adopts a less compact conformation in this phase, with several chromosome segments looping out from the mesh-like structure that is less clearly defined than in the other conditions. This effect was quantified by measuring the radius of gyration (*R_G_*) of the polymers across the ensemble,

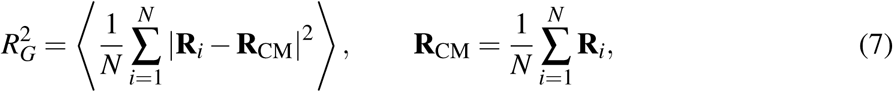

which shows that anaphase structures are significantly more extended than those in all other conditions (Fig. 2D). Inspection of individual chromosomes suggests that this increased extension may reflect a state in which centromeres have already been pulled toward the spindle pole bodies in the mother and bud, while parts of the chromosome arms remain extended toward the opposite side. The resulting reduction in contacts between these two compartments would account for the observed decompaction. At later stages of anaphase, condensin would then compact chromosome arms toward the centromeres through loop extrusion, thereby completing chromosome segregation [35, 61–64]. Since the Hi-C map considered here corresponds to an earlier stage, the inferred structures are expected to retain this extended arm configuration across the two nascent nuclei.

The G0 simulations showed a larger *R_G_* than the G1 simulations, despite the smaller nuclei and greater chromatin compaction observed in quiescent cells [34, 54, 65, 66]. Hi-C-derived structures have a relative spatial scale, so their conversion to physical distances requires calibration for each condition. Calibration against experimental inter-locus distances gave σ_G1_ = 74 nm (Fig. 1F).

Because corresponding G0 measurements are unavailable, the measured nuclear radii serve as a proxy for the ratio of chromosomal radii of gyration, rather than as direct measurements of *R_G_*. Using simulated values of *R_G_*_,G0_ = 6.7σ and *R_G_*_,G1_ = 6.6σ, and experimental nuclear radii of 1.5 µm and 1.7 µm, respectively [34], gives

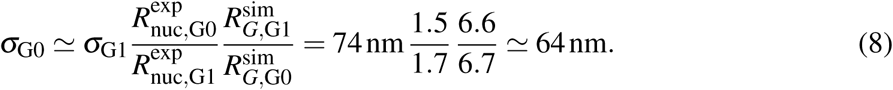

This condition-specific calculation accommodates the smaller nuclear size in G0 and is consistent with the greater chromatin compaction expected from the changes in transcription and global histone deacetylation observed in quiescence [65–68]. Inter-locus distance measurements across the cell cycle would allow direct phase-specific calibration and improve the model’s quantitative predictions.

### 3.3 Emergent features appear upon FI-Chrom Modeling

We next examined centromere and telomere clustering in G1, anaphase and G0 (Fig. 3A–C). The intermediate conditions, Mid-S, S/G2 and G2/M, are presented in Supplementary Fig. S7. Compared to the full three-dimensional bead distribution of the genome, centromeres in both G1 and anaphase were preferentially located within a distinct compartment (Fig. 3B). As such, we were able to recover centromere clustering without explicitly including the SPB as in the YSM. The extent of clustering was quantified by measuring the *R_G_*density of the centromeres beads, and comparing it to the *R_G_*of the Intra-D and Inter-D models (distribution in Fig. 3F). For these models, for which centromeres don’t carry any special identity, their *R_G_* is comparable to the *R_G_* of the whole system meaning that centromeres are uniformly distributed across the full volume occupied by the beads. For G1, anaphase and G0, the density of *R_G_*centers at lower values. Reflecting that centromeres are concentrated in their own compartments. Across the considered conditions, centromeres in anaphase result comparatively more clustered then in G1 and G0. These differences are consistent with evidence that the spatial extent of centromere clusters varies during the cell cycle, with an increased compaction observed in anaphase by super-resolution microscopy [69]. They may reflect changes in the length and organization of kinetochore microtubules or in kinetochore–microtubule attachments [70, 71]. In particular, microscopy has shown that, during quiescence, kinetochores remain attached to microtubules but redistribute along an elongated nuclear microtubule array, consistent with the greater centromere dispersion inferred here for G0 [71].

**Figure 3:**
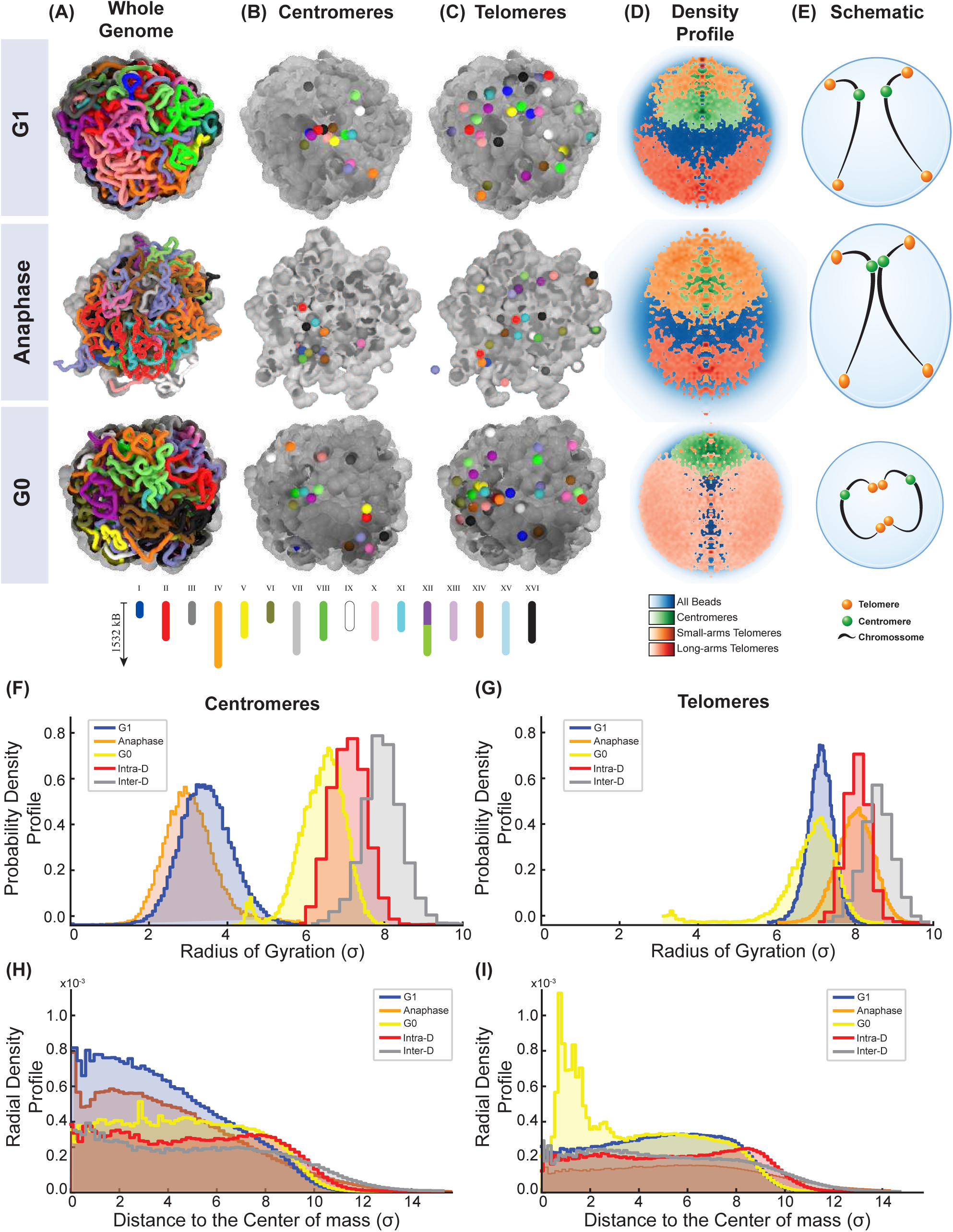
Emergent structural features of the FI-Chrom model. (A–C) Representative whole-genome conformations (A), centromere positions (B), and telomere positions (C) in G1, anaphase, and G0. (D) Spatial density profiles of centromeres and telomeres on short (< 200 kb) and long chromosome arms, aligned along the axis connecting the genome and centromere centers of mass. (E) Schematic summary of centromere and telomere positioning. (F, G) Probability distributions of the radius of gyration of centromeres (F) and telomeres (G), quantifying their spatial clustering. (H, I) Radial distributions of centromeres (H) and telomeres (I), measured from the genome center of mass.

A key difference between this model and the YSM, is that in the YSM centromeres are permanently tethered to the SPB, whereas in our model the centromere compartment is defined dynamically by the mutual affinity between each other. As such, individual centromeres can transiently detach from the region of high centromere density and subsequently return to it. The dynamical stability of the SPB has been quantified by fluorescent tagging of an SPB component and a fluorescent operator tagged near the centromere, and their distance has been measured with high resolution in living cells [72]. Microtubules grow and shrink continuously, and the SPB and centromere were never closer than ∼ 200 nm in G1 phase [15, 72]. This dynamical behavior might seldomly result in lower Hi-C scores then what would be expected by direct interactions. Our results may also reflect that Hi-C does not capture N-body interactions, and this affects the precision of our model when considering the coincidential association of a large number of centromeres. Such interactions could, in principle, be incorporated into our framework to improve this aspect of the model. That would require the integration of multicontact genome-mapping approaches such as SPRITE [73] or *in vivo* imaging data. However, a dataset of this type focused on budding yeast centromeres under all the conditions considered here is still unavailable in literature.

Alternatively, this result may indicate that the dynamical stability of SPB association should itself be reconsidered. In support of this possibility, disruption of the kinetochore–microtubule link leads to a perturbation of centromere clustering [70]. And individual centromere detachment from the SPB has been recently observed in yeast following DNA damage by Haber and collaborators [74], and is dependent on Smc5/6 binding to microtubules [75], challenging the notion that the SPB association of all centromeres is completely stable. This hypothesis should also be evaluated in light of recent observations indicating that centromere clustering is a widespread nuclear organization strategy adopted by cells from diverse species across the eukaryotic evolutionary tree, including human haploid cells defective for the condensin subunit SMC2 [76]. Importantly, centromere clustering does not always give rise to a single nuclear centromere domain at this level. For example, Chavan *et al.* quantified the D1 centromere-satellites-associate protein in *Drosophila melanogaster* male and female germline stem cells, supporting the view that chromocenter-associated heterochromatin in the germline can be organized into multiple nuclear foci rather than a single domain [77]. Thus, signatures of centromere clustering in contact maps should not be taken as evidence for a complete, or canonical, Rabl conformation, nor should they be confused with the formation of a single centromere cluster within the nucleus. Taken together, based on our simulations, we can only conclude that the Hi-C maps of budding yeast, taken alone, are inconclusive in this regards. And this aspect of chromatin organisation deserves further investigation.

The model predicts a complementary pattern for telomeres, with stronger clustering in G0 than in G1 and anaphase (Fig. 3C, and Fig. 3G). This agrees with the formation of a telomere hypercluster in G0, which is absent during other growth phases [34, 56, 57]. To determine how these changes in clustering reshape nuclear architecture, the radial localization of centromeres and telomeres was subsequently examined. As discussed in the Introduction, the nuclear organization of budding yeast is symmetric around the axis connecting the SPB and the nucleolus. Since Hi-C data do not capture the nucleolus itself, in our simualtions chromosome XII was interrupted at the rDNA repeat region, allowing the two flanking segments to fluctuate independently. In this way, the space between them effectively represents the missing connectivity associated with the nucleolar region. In order to capture the symmetry in the Rabl configuration, we decided instead to align all structures in the cylindrical coordinate system defined by the axis connecting the center of mass of the centromeres to the center of mass of the whole system. This allowed us to estimate the density in the (*z*, *r*) coordinates for each bead (See SI for details).

Similarly to what has been done from imaging data [14, 15], we plot the density of centromeres and telomeres, separating telomeres connected to short chromosomal arms (< 200 kb) from telomeres connected to long chromosomal arms (Fig. 3D). Strikingly, we find similar patterns as the ones reported in the imaging data. Centromeres are clustered at the periphery towards the center of the nucleus. In both G1 and anaphase, telomeres preferentially occupy the periphery of the nucleus [21, 78], with small chromosomal arms’ telomeres located closer to centromeres than longer chromosomal arms’ [14]. In anaphase, the mean structure is elongated on the axis that connect the centromeres to the telomeres of the longer chromosomal arms, that is an hallmark of the symmetry breaking expected from chromosomes being segregated apart by the microtubules. In the G0 phase, telomeres detach from the nuclear envelope and occupy the full volume of the nucleus (Fig. 3E).

Association with the nuclear envelope was quantified by measuring the radial density profiles of centromeres, Fig. 3H) and telomeres (Fig. 3I). This analysis support a global distribution of centromeres farther from the nuclear envelope in G1 and anaphase than in G0. Microscopy shows that in G0 centromeres redistribute along an elongated microtubule array spanning the nucleus, including at its distal tip [71], potentially bringing a subset closer to the nuclear envelope. In contrast, telomeres are substantially more enriched near the nuclear center in G0 than in either anaphase or G1. This result is particularly unexpected because it challenges the prevailing view that contact maps contain little or no information about positional features such as association with the nuclear membrane. This view is supported by the observation that contact maps obtained from wild-type thymocytes and Lamin B receptor-deficient thymocytes are highly similar, despite the fact that the latter exhibit an inverted nuclear architecture by fluorescence microscopy, with heterochromatin largely displaced away from the nuclear envelope [79]. In contrast, our results show that yeast Hi-C contact maps retain sufficient information to infer the spatial positioning of telomere clusters, and that this information can be recovered by our machine-learning approach. Finally, our simulations support a close relationship between nuclear-envelope anchoring and telomere hypercluster formation. Telomeres are split in multiple loci when associated to the nuclear envelope in exponential growth [80], while they are associated to an hypercluster in the quiescent phase [34]. Our result is consistent with recent work showing that Esc1-mediated anchoring to the nuclear envelope is a major antagonizer of telomere clustering, and that its progressive loss of function upon entry into quiescence promotes telomere hyper-cluster formation [25].

### 3.4 Dynamic analysis of chromosome components

We then leveraged the ability of our model not only to generate an ensemble of conformations consistent with the contact maps, but also to characterize their dynamics and fluctuations over time. Because FI-Chrom is trained on static Hi-C contact probabilities, the resulting trajectories should not be interpreted as a direct reconstruction of the real temporal history of individual chromosomes in living cells. Instead, they represent the effective polymer dynamics associated with an equilibrium ensemble constrained by the experimental Hi-C maps. This provides a useful test of whether the structural information inferred from Hi-C is compatible with independently measured chromatin mobility.

For each genomic bead, the mean-square displacement (MSD) as a function of the lag time was computed,

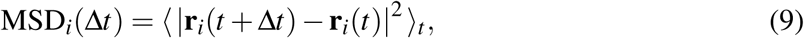

where r*_i_*(*t*) is the position of bead *i* at time *t*, and the average is performed over time origins and independent replicas. Spatial units were converted to physical units using the calibration obtained from the comparison with experimental inter-loci distances in G1. The simulation time scale was then chosen by comparing the simulated MSD with live-cell tracking measurements for the corresponding chromosomal locus.

Focusing first on the G1 condition, for which quantitative experimental measurements of chromatin mobility are available, the simulated MSD of a locus located on chromosome IV, corresponding to locus 181 at approximately 430 kb, was compared with the experimental measurements reported by Arbona *et al.* [23, 81]. After this time-scale calibration, the FI-Chrom trajectory reproduces the scaling of the experimental MSD over the measured time window (Fig. 4A). Importantly, these dynamical data were not used during the optimization procedure. Thus, the agreement with experimental mobility emerges from the combination of the Hi-C-inferred effective force field and the generic polymer dynamics used to sample the model.

**Figure 4:**
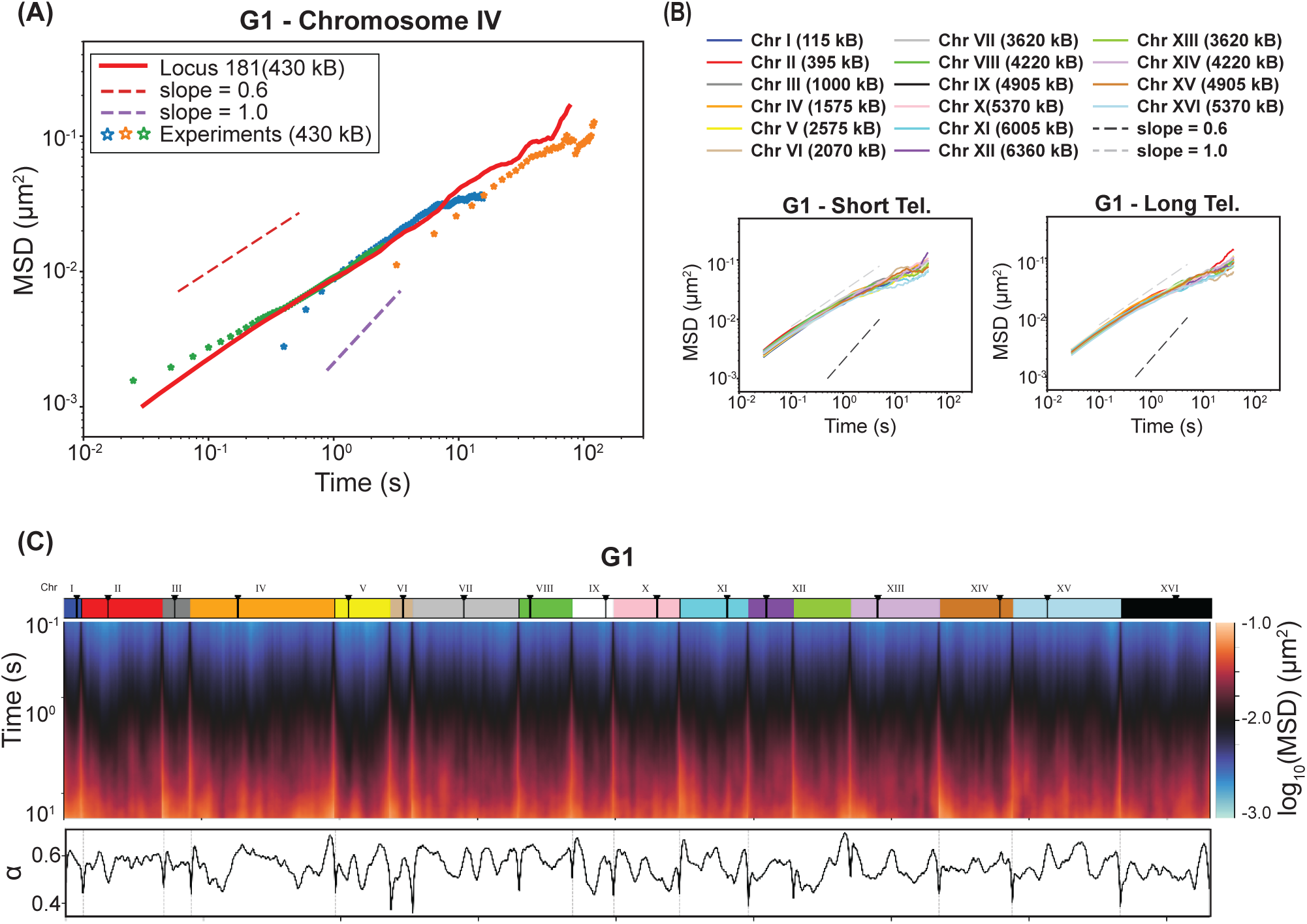
Analysis of the chromatin dynamics. Chromatin dynamics in the G1 FI-Chrom model. (A) Mean-square displacement (MSD) of a locus at approximately 430 kb on chromosome IV, comparing simulations (solid line) with experimental measurements (stars) [23, 81]. Dashed lines indicate reference scaling exponents of 0.6 and 1.0. (B) MSD profiles of telomeres on short and long chromosome arms. (C) Genome-wide heatmap of log_10_(MSD) as a function of genomic position and lag time. Centromere positions are indicated above the heatmap. The lower panel shows the fitted exponent α for each 5-kb bead, defined by MSD(Δ*t*) ∼ Δ*t*^α^ .

The MSD displays a clear subdiffusive behavior, with an effective scaling exponent below the value expected for normal diffusion. This is consistent with previous experimental and theoretical studies showing that yeast chromatin dynamics are largely compatible with a Rouse-like polymer regime, modulated by transient locus-specific interactions [16, 17]. In this regime, the motion of a chromatin segment is constrained by its connectivity to neighboring genomic regions and by the collective relaxation of the chromosome polymer. The fact that our model recovers this behavior suggests that the inferred three-dimensional organization does not impose an unrealistic dynamical constraint on the chromatin fiber.

Next, the model was used to compare the mobility of chromosome components with distinct nuclear organization patterns. In G1, telomeres are preferentially associated with the nuclear periphery, while centromeres form a compact compartment associated with the Rabl-like organization of the yeast genome. The MSD of telomeric loci on short and long chromosome arms was then quantified (Fig. 4B).

The MSD was then calculated for every 5-kb bead and plotted against genomic position and lag time (Fig. 4C). This genome-wide map reveals that chromatin mobility is not uniform along chromosomes. Regions close to centromeres generally display reduced motion, consistent with their participation in the centromere cluster. In contrast, chromosome arms exhibit broader variations in mobility, reflecting differences in local polymer environment, chromosome length, and long-range interactions learned from the Hi-C map. This analysis reveals that telomere mobility is heterogeneous across chromosomes. Part of this heterogeneity is expected from polymer connectivity: telomeres belonging to short chromosome arms are more strongly influenced by the proximity of the centromere compartment, whereas telomeres on long arms can explore a larger effective conformational space. Thus, even in the absence of explicit nuclear-envelope tethers, the Hi-C-inferred interaction landscape is sufficient to generate component-specific dynamical differences.

The local dynamical regime, quantified by the MSD of each bead, was then fitted to a power law, MSD(Δ*t*) ∼ Δ*t*^α^, where α is the anomalous diffusion exponent. Across the genome, most loci show α < 1, confirming that the simulated yeast genome remains in a subdiffusive regime.

This result is consistent with the interpretation of chromatin as a connected polymer embedded in a crowded nuclear environment, where local motion is constrained by both chain connectivity and collective chromosome organization.

The same bead-resolved MSD analysis was performed for all other cell-cycle stages, and the corresponding heatmaps are reported in the Supplementary Information (Supplementary Fig. S8–S12). Overall, these profiles did not reveal substantial stage-dependent changes in the spatial pattern or scaling regime of chromatin mobility. Therefore, we focus the main-text discussion on the G1 condition, where direct comparison with experimental live-cell tracking data is available.

## 4 Discussion and conclusion

In this work, we introduce a data-driven polymer model of the entire *Saccharomyces cerevisiae* genome throughout the cell cycle. Using our improved method based on FI-Chrom and Serpentine [12, 37], we infer an effective force field directly from experimental Hi-C contact maps, without explicitly imposing canonical features of nuclear architecture, such as centromere clustering, telomere positioning, or chromosome territoriality, as is done in the YSM [16, 17, 23, 24]. From only chromosome lengths, polymer connectivity, and measured contact probabilities, the model generates three-dimensional ensembles of genome conformations in G1, mid-S, S/G2, G2/M, anaphase, and G0. Remarkably, these minimal data-driven ingredients are sufficient to recapitulate the major hallmarks of yeast nuclear organization and their remodeling across the cell cycle: the characteristic radial distributions of chromosomes, centromere and telomere clustering, the striking elongation of long chromosome arms during anaphase, and the release of telomeres from the nuclear envelope upon entry into G0. Thus, complex, phase-specific features of genome architecture emerge directly from the experimental contact maps rather than from prescribed geometrical constraints.

The YSM model has been highly successful in explaining the spatial organization of the yeast genome in G1 and in providing a mechanistic basis for several experimental observations. Our work takes an alternative route. By removing most of these explicit biological assumptions, we asked how much of yeast nuclear organization can be recovered from Hi-C data by itself. The answer is not trivial, because Hi-C measures population-averaged contact frequencies rather than absolute three-dimensional positions or single-cell trajectories. Nevertheless, our results show that the contact maps contain sufficient information to recover several known features of yeast genome organization and to predict additional structural and dynamical properties.

In G1, the FI-Chrom model reproduced chromosome boundaries, centromere-centromere contacts, the mating-type locus interaction on chromosome III, and the scaling of contact probability with genomic distance. The model also generated three-dimensional conformations that were compatible with independent imaging measurements of inter-loci distances. Applying this strategy across the cell cycle revealed progressively stronger chromosome territoriality from G1 to G2/M, consistent with cohesin-dependent organization and reduced chromosome intermixing. By contrast, anaphase showed a globally decompacted, extended, and asymmetric genome configuration, with an increased radius of gyration. This organization is consistent with centromeres being pulled toward opposite spindle pole bodies while the long chromosome arms remain stretched, recapitulating the polarized architecture of chromosome segregation.

Several nuclear landmarks emerge from the inferred interaction landscape without being explicitly imposed. In particular, centromeres form a compact compartment in G1 and anaphase, even though the model does not include the spindle pole body or microtubules. This compartment becomes more compact in anaphase [69]. These results suggest that pairwise contact information is sufficient to reproduce a centromere-enriched domain characteristic of the Rabl-like organization of budding yeast. However, because FI-Chrom relies on pairwise interactions, it cannot distinguish between true many-body tethering and an effective mutual attraction between centromeric regions. Thus, while the model recovers the contact-map signature of centromere clustering, it does not establish the underlying physical mechanism. Higher-order contact measurements or simultaneous live-cell imaging of multiple centromeres would be required to resolve this point.

Telomeres displayed a complementary behavior. In G1 and anaphase, telomeres preferentially occupy peripheral regions, consistent with their association with the nuclear envelope during proliferative growth. In G0, the model predicts a redistribution of telomeres toward the nuclear interior and increased telomere clustering. This is consistent with the dynamic of the telomere hypercluster formation observed in quiescent cells and supports the idea that loss of nuclear-envelope anchoring promotes telomere coalescence [25]. Thus, even in the absence of explicit telomere tethers, the Hi-C-inferred effective force field captures the transition between peripheral telomere organization in cycling cells and telomere clustering in quiescence.

The polymer trajectories reproduce the experimentally observed subdiffusive motion of a chromatin locus in G1 and predict heterogeneous mobility across the genome, with centromeres generally moving less than chromosome arms. Bead-resolved MSD profiles for the other stages (Supplementary Information) show no major change in the global dynamical regime, suggesting that the subdiffusive motion characteristic of a polymer persists throughout the cell cycle. Thus, chromosome territoriality, centromere clustering, and telomere positioning can be extensively reorganized without large changes in global chromatin mobility. Hi-C and MSD therefore provide complementary information: Hi-C reports relative contact frequencies, whereas MSD measures locus mobility constrained by the inferred physical environment.

The predicted genome-wide heterogeneity in mobility and the distinct behavior of telomeres therefore provide independent, experimentally testable outputs. In particular, telomere mobility depends on chromosome-arm length and the stage-specific interaction landscape. These predictions can be tested directly through live-cell tracking.

The model has two main limitations. First, it infers an effective equilibrium ensemble independently for each cell-cycle stage and therefore does not describe temporal transitions between stages. Active processes, including DNA replication, cohesin loading, condensin-mediated loop formation, microtubule pulling, telomere anchoring, and nucleolar remodeling are not modeled explicitly; their effects are absorbed into stage-specific force fields. Thus, the framework is not a mechanistic model of cell-cycle progression. Future extensions could explicitly incorporate these processes. Second, the model assumes a homogeneous population within each stage, implying that cell-cycle position is the main source of conformational variation in budding yeast. This assumption may not hold in organisms with more complex differentiation programs. More generally, population-averaged measurements such as Hi-C cannot readily distinguish strong effects confined to small subpopulations from weaker effects shared by all cells. In the future, combining machine learning with polymer-physics approaches may help identify such heterogeneity, including bimodal conformational states.

Despite these limitations, the approach provides a flexible platform for generating quantitative, experimentally testable predictions. The most direct validation would come from fluorescence imaging at the same cell-cycle stages and in the quiescent condition modeled here. In particular, measurements of intra-arm distances, centromere and telomere positions, and locus-specific MSDs along replication, in G2/M, anaphase, and G0 would support condition-specific calibration of spatial and temporal units. The integration of such data would enable quantitative testing of the predicted anaphase decompaction, telomere redistribution in G0, and relative stability of centromere clustering across the cell cycle, while guiding further refinement of the model. Further, it is possible to integrate the result of our parameter optimization with structural elements introduced ad-hoc, this can be done both before optimization to reduce and optimize the parameter space as well as after optimization to test the effect of perturbations to chromatin structure.

In conclusion, we present the first machine-learning-based inversion of whole-genome Hi-C maps across the complete cell cycle of a eukaryotic microorganism. By transforming contact data into three-dimensional chromosome molecular dynamics simulations, our framework reveals major features of genome organization that were neither prescribed nor directly apparent from the maps alone. More broadly, it establishes a scalable route from Hi-C data to data-driven physical models of chromatin architecture, opening new opportunities to resolve population heterogeneity, incorporate higher-order interactions, and investigate the larger and more complex genomes of multicellular organisms.

## Acknowledgments

The authors acknowledge helpful discussions with Jean-Marc Victor and Angela Taddei. This research was conceived at the Genome Architecture and Function Summer School and Workshop 2023, in Sofia (Bulgaria), the authors acknowledge the support of the International Physics of Living Systems network for organizing and supporting this conference, sponsored by CNRS (France), NSF (USA) and NSF (Bulgaria). This research was supported by the Center for Theoretical Biological Physics, sponsored by the NSF (Grants PHY-2019745, PHY-2607619, PHY-2014141, and PHY-2210291) and by the Welch Foundation (Grant C-1792). JNO is a Cancer Prevention and Research Institute of Texas (CPRIT) Scholar in Cancer Research. VGC acknowledges support from the NSF Award PHY-2609969 and the NVIDIA Academic Grant Program. VFS and JNO acknowledge the support of the Paris Sciences et Lettres university PSL Global Seed Fund and Rice University to foster the collaboration. ABOJ acknowledges the Robert A. Welch Postdoctoral Fellow Program. We would like to thank AMD (Advanced Micro Devices, Inc.) for donating critical hardware and support resources from its HPC Fund, which made this work possible.

## Supplementary Information

### Theory

#### Inverse Statistical Mechanics and the Maximum Entropy Approach

The maximum entropy (MaxEnt) principle [1, 2] is a method used to infer the least biased probability distribution given a set of constraints. In the context of chromosome simulations, the MaxEnt approach is applied to generate a bias potential that incorporates experimental data without introducing unnecessary assumptions [3]. The bias potential is constructed in such a way that the resulting distribution of configurations is consistent with known experimental observables (e.g., pairwise distances and contacts between chromosome regions).

The MaxEnt approach ensures that we only incorporate the necessary information to satisfy the imposed constraints while maximizing the entropy of the system, thus maintaining the most unbiased distribution possible [4]. This makes it particularly suitable for systems like chromosomes, where the complexity and variability in configuration space require minimal assumptions.

Starting from the potential describing a homopolymer *U*_HP_(r), whose explicit expression is provided in the following sections; For the homopolymer potential and for a generic phase space observable function *φ* (r), a simulation in the canonical ensemble yields the following value:

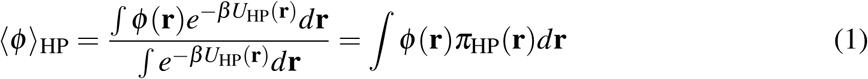

where r is the vector characterizing the positions in Cartesian space of all the loci in the chromosome, *π*_HP_(r) is the probability density for the homopolymer model, and 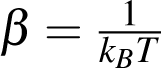

Following the Maximum Entropy approach, we consider the probability density *π*_ME_(r) that reproduces the experimental values of a certain set of *n* observables *φ_i_*(r). The following constraint equations define such a probability density:

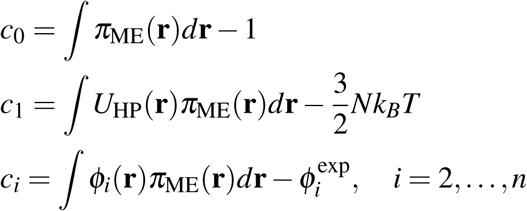

Each of the constraint equations must be equal to zero; the first equation ensures that *π*_ME_(r) is normalized, the second fixes the average potential energy to be equal to the thermal energy 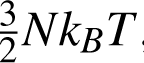, and the last *n* equations ensure that the expectation values and the experimental values coincide for each one of the considered observables.

To determine the probability density *π*_ME_(r), we maximize the information entropy:

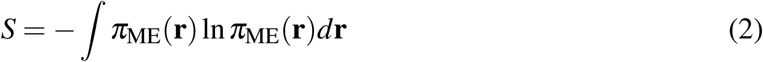

subject to the constraints. This approach is based on the fact that a constrained maximization of the entropy is equivalent to minimizing the amount of additional information built into the distribution other than the one contained in the constraints themselves.

Using Lagrange multipliers, we obtain the following extremum condition:

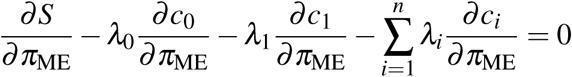

which leads to the probability distribution:

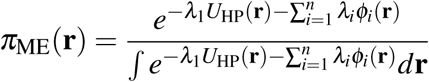

Recognizing that *λ*_1_ coincides with *β* , we can think of the maximum entropy probability distribution *π*_ME_(r) as the distribution sampled from the maximum entropy potential energy:

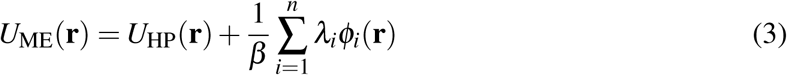

In our case, *φ_i_*(r) measures the contact probability of each genomic pair formed between any two loci separated at a distance r.

The *λ_i_*are the Lagrangian multipliers whose values can be obtained via minimization of the objective function Γ(*λ*) defined as

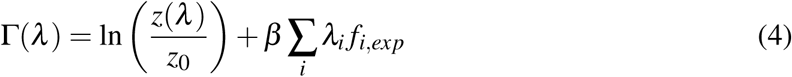

where *β* = 1/*k_B_T* , with *k_B_* as the Boltzmann constant and T as the temperature. (*λ*) and *z*_0_ are the partition functions for the homopolymer with and without the maximum entropy correction.

Following the approach by Zhang & Wolynes [3], we simplify the objective function using the cumulant expansion. However, here, we will use only the first term of the cumulant; the simplified objective function becomes:

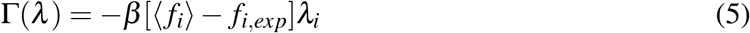

Although there is no guarantee that the resulting *λ* ’s will exactly reproduce the data, we recognize that Eq. 5 provides only an approximate solution for *λ* . Therefore, when ⟨ *f_i_*⟩ does not match *f_i_*_,*exp*_, we apply the Adam Optimization algorithm to refine these values, aiming for eventual convergence with the experimental measurements. The details of Adam Optimization will be explained in the next section.

#### Adam Optimization

The optimization procedure plays a crucial role in determining the best values for the Lagrange multipliers *λ_i_*, which represent the interaction strength between loci pairs *i* and *j*. Given a polymer consisting of *N* beads, the number of interaction parameters is *S* = 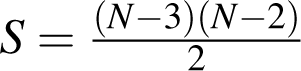. This formula excludes the interactions between the first and second nearest neighbors, as these are already considered in the bond and angle potentials. For example, in the case of human chromosome 1, modeled at a resolution of 50 kb with *N* = 4960 beads, there are *S* = 12, 288, 403 interaction parameters to be optimized. The number of parameters grows quadratically with the number of beads, making it necessary to employ an efficient optimization algorithm for handling such large parameter spaces. To optimize the millions of parameters in the FI-Chrom model, we employ the Adam optimizer [5], a widely used algorithm in machine learning that adapts the learning rate for each parameter based on estimates of the first and second moments of the gradients. This method is particularly well-suited for high-dimensional parameter spaces, such as those encountered in chromatin structure modeling.

Adam is based on adaptive estimates of lower-order moments (mean and variance) of the gradients. Specifically, at each iteration *t*, the gradient of the loss function with respect to the parameters, *g_t_*, is used to update two estimates:

- The first moment, which is an exponentially weighted moving average of the gradient, denoted by *m_t_*.
- The second moment, which is an exponentially weighted moving average of the squared gradient, is denoted by *v_t_*.

These are computed using the following recursive relations:

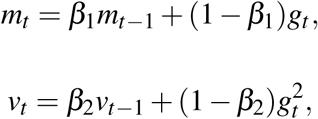

where *g_t_*is the gradient of the loss function at iteration *t*, and *β*_1_= 0.9, *β*_2_ = 0.999 are decay rates that control how quickly the moving averages adjust to the incoming gradients.

To counteract the bias introduced in the initial steps of the optimization process, Adam employs bias correction terms for both *m_t_* and *v_t_*, given by:

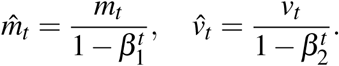

The updated parameters, *θ_t_*, are computed by adjusting the previous parameters, *θ_t_*_−1_, in the direction of the negative gradient, with an adaptive step size based on the corrected first and second moments:

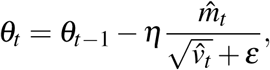

where *η* is the learning rate, and *ε* is a small constant added for numerical stability (typically *ε* = 10^−8^).

In the context of the FI-Chrom model, the gradient *g_t_* is calculated based on the difference between the experimental and simulated contact probabilities:

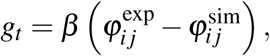

where *β* = *k_b_T*, combining the Boltzmann constant *k_b_* and the information-theoretic temperature *T* , and 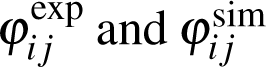 denote the experimental and simulated contact probabilities, respectively, for loci pair (*i*, *j*). The gradient quantifies the discrepancy between the two probabilities, driving the optimization process to minimize the difference between experimental data and the simulated model at each iteration.

By iteratively updating the interaction parameters using the Adam algorithm, we ensure efficient convergence towards an optimal set of parameters, even in cases where the parameter space is large and complex. The adaptive learning rates for each parameter allow for a fine-tuned optimization process that balances exploration and convergence speed, which is particularly important for models that involve millions of interaction parameters, such as high-resolution chromatin models.

#### Homopolymer Energy Function

The homopolymer potential models a generic bead-spring polymer in which each bead represents a genomic segment of 5 *kb* in sequence. The potential energy *U_HP_*(⃗*r*) describes a spatially confined self-avoiding polymer (no nonbonding attractive term is included here) and serves as a support for the features added by using the maximum entropy principle.

This potential consists of the following five terms, *U*_FENE_ , *U*_Angle_ , *U_hc_*, *U_sc_* and *U_c_*:

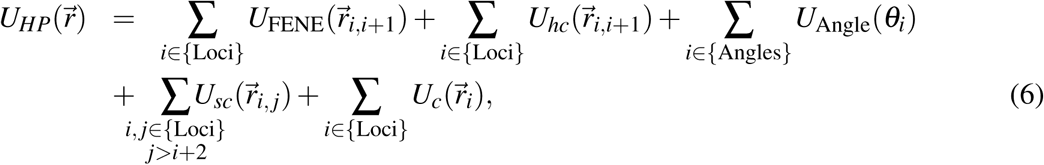

where,

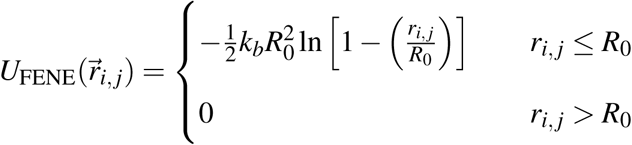

*U*_FENE_ (Finite Extensible Nonlinear Elastic potential) is the bonding term applied between two consecutive monomers.

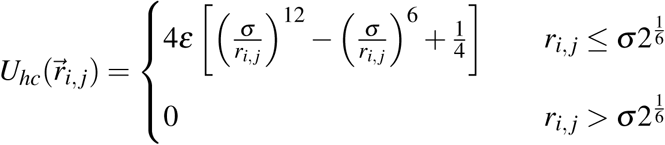

*U_hc_*(⃗*r_i_*, *_j_*) is the hard-core repulsive potential, included to avoid overlap between bonded monomers.

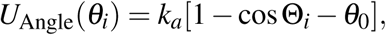

a three-body potential applied to all connected three consecutive monomers, where *θ_i_*is the angle defined by two vectors⃗*r_i_*,*_i_*_+1_ and⃗*r_i_*,*_i_*_−1_.

The non-bonded pairs is defined by a soft-core repulsive interaction in the following form:

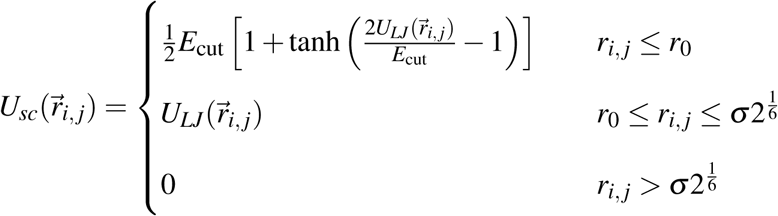

The expression correspond to the Lennard-Jones potential *U_LJ_*(⃗*r_i_*, *_j_*) = 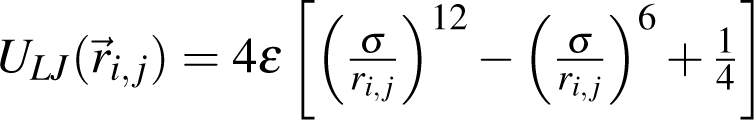 capped off at a finite distance, thus allowing for chain crossing at finite energetic cost. *r*_0_ is chosen as the distance at which *U_LJ_*(⃗*r_i_*, *_j_*) =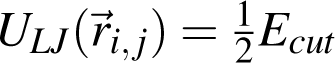

The confinement potential *U_c_* describes the interaction between the chromosome and a spherical wall, whose size is chosen to enforce a volume fraction of 0.1; which is consistent with the experimentally determined density of chromatin. The spherical wall is included to mimic the interaction with the nucleus envelope that chromosomes experience inside the cell. Each monomer *i* of the chromosome interacts with its nearest point on the wall *r_np_* through the potential *U_hc_*(*r_i_*_,*np*_).

### Hi-C data and normalization procedure

To investigate cell-cycle-dependent changes in genome organization, we trained and analyzed six independent budding yeast Hi-C datasets corresponding to G1, Mid-S, S/G2, G2/M, Anaphase, and G0 (starvation), obtained from the published dataset generated by Romain Koszul’s laboratory [6]. For each cell-cycle stage, an independent FI-Chrom force field was inferred from the corresponding experimental contact map, allowing stage-specific changes in both intraand inter-chromosomal interactions to be captured.

In addition to the FI-Chrom optimized models, we considered two reference polymer models, denoted here as the intra-dominant and inter-dominant models. These models were used as theoretical baselines to assess the structural effects introduced by the inferred non-bonded interactions and to determine the extent to which generic polymer physics alone can reproduce the experimentally observed cell-cycle-dependent Hi-C patterns.

Experimental contact maps were constructed from raw ligation counts at 5 kb resolution. The raw contact matrices were balanced using the Knight–Ruiz (KR) normalization procedure, which rescales the matrix such that all rows and columns have equal sums. Let the resulting balanced contact matrix be denoted by *C*.

To obtain the experimental contact probability matrix, *P*_Hi−C_, each row *i* of the balanced matrix was rescaled by its maximum contact value:

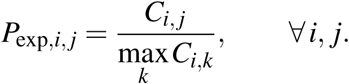

Thus, the strongest contact associated with each genomic locus is assigned a normalized value of 1, while all other contacts are expressed relative to this maximum. In practice, the largest contact frequency is generally associated with self-contacts or contacts between neighboring genomic bins, reflecting the strong local connectivity expected for a beads-on-a-string polymer representation. This normalization therefore maps the experimental Hi-C signal onto the range 0 ≤ *P*_exp,*i*,_ *_j_* ≤ 1, providing the target contact values used in the FI-Chrom optimization procedure.

### Analysis Tools

#### Radial Density Profile

The radial density profile can be defined as:

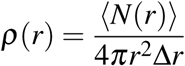

where *r* is the distance from a genomic locus to the centroid, *N*(*r*) is the number of genomic loci found in the spherical shell *r*, *r* + Δ*r* , and Δ*r* = *R*_0_/200 is the size of the numerical grid. *R*_0_ is the size of the confinement to ensure a density of 0.1, and is chromosome dependent. ⟨•⟩ indicates averaging over the ensemble of chromosome conformations. When calculating the density profile for a given chromatin type, only loci of that type are included in counting *N*(*r*). To compare different chromatin types and loci with different gene expressions, we normalize the radial profiles by the number of loci of a given type *N* Type.

## Supplementary Results

**Figure S 1:**
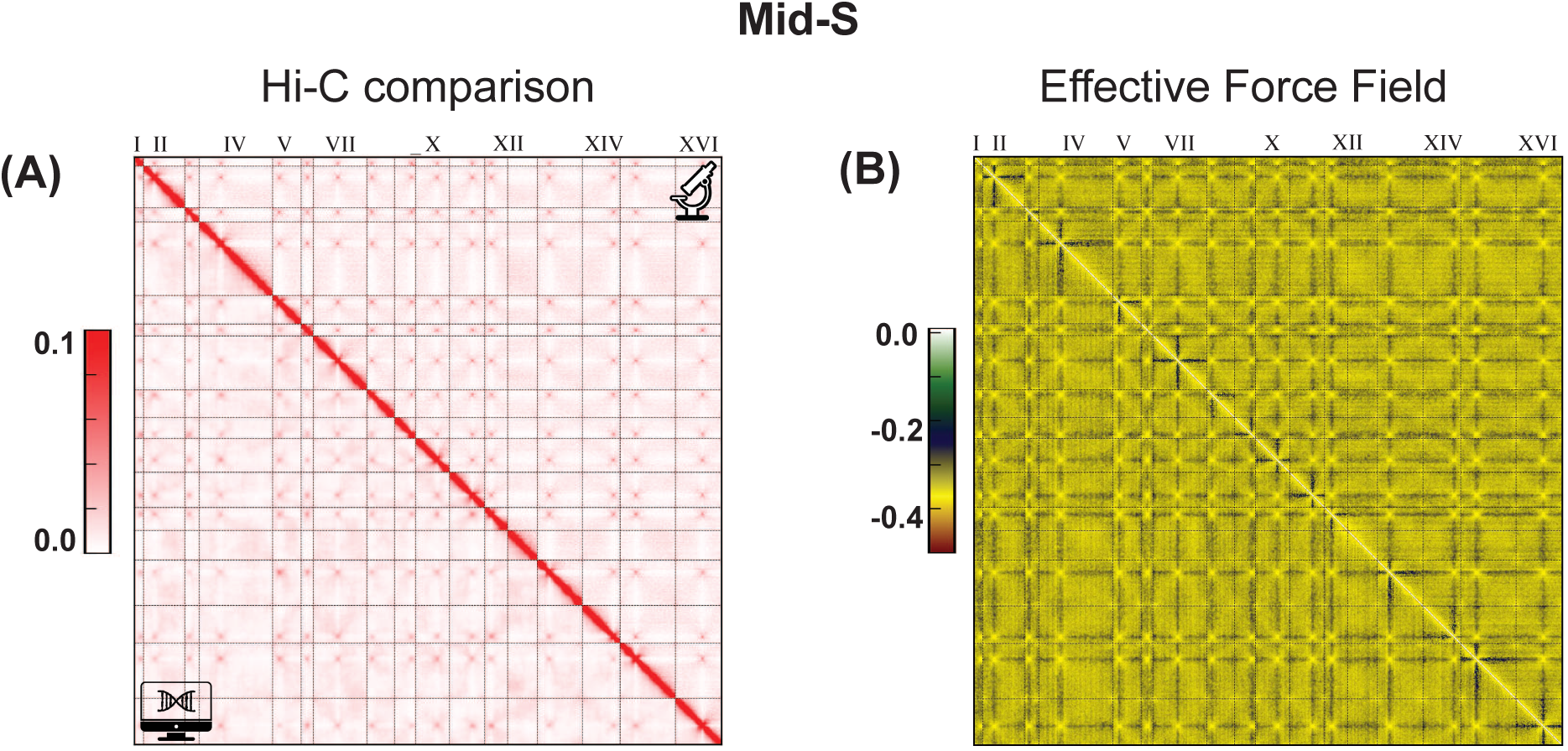
A) Comparison of Hi-C (upper triangle) and simulation (lower triangle) contact maps for Mid-S phase. (B) Trained EFF for the Mid-S phase obtained using the FI-Chrom method [7] (see Methods for more details). Pairwise energy parameters (*λ*) range from −0.4 (attractive) to 0.4 (repulsive).

**Figure S 2:**
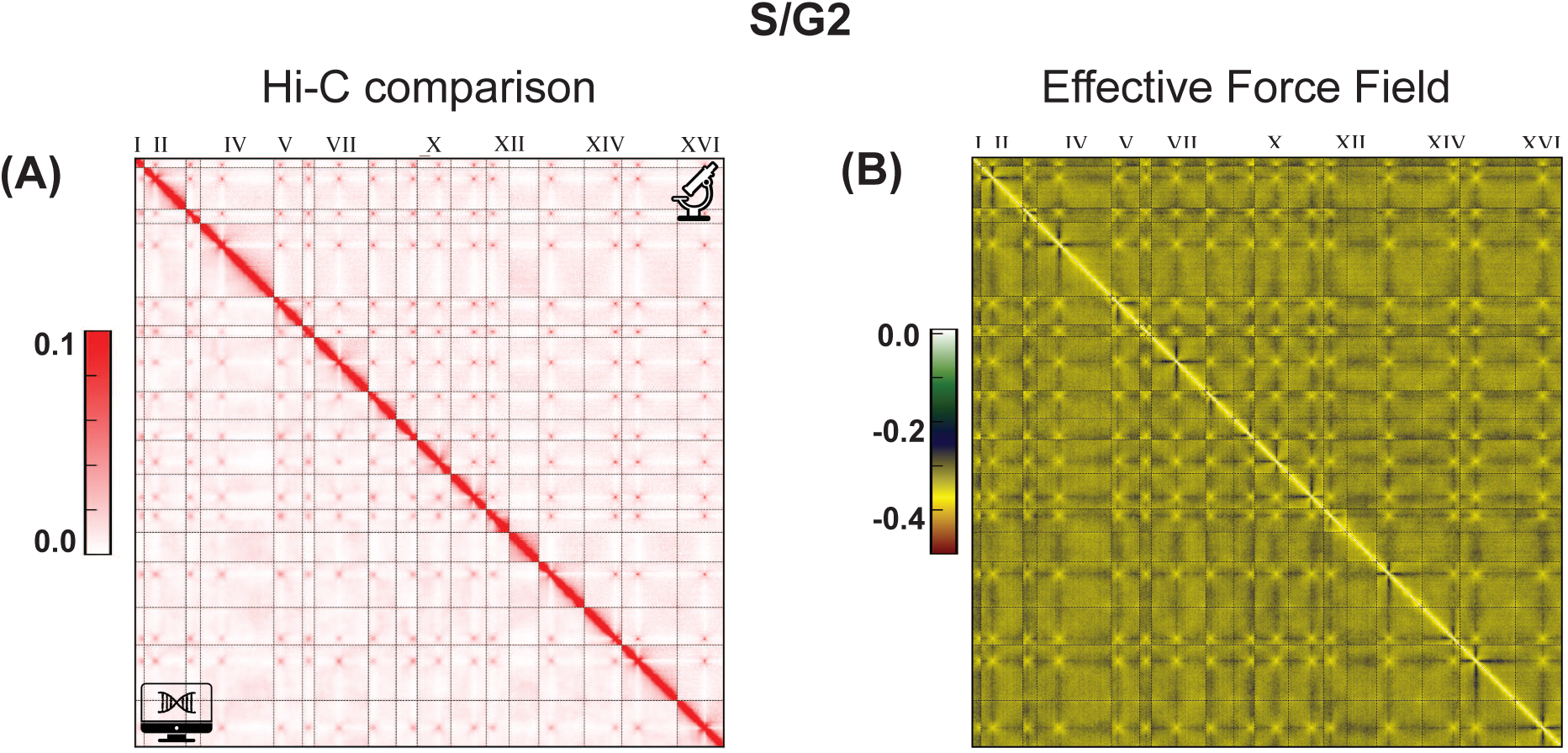
A) Comparison of Hi-C (upper triangle) and simulation (lower triangle) contact maps for S/G2 phase. (B) Trained EFF for the S/G2 phase obtained using the FI-Chrom method [7] (see Methods for more details). Pairwise energy parameters (*λ*) range from −0.4 (attractive) to 0.4 (repulsive).

**Figure S 3:**
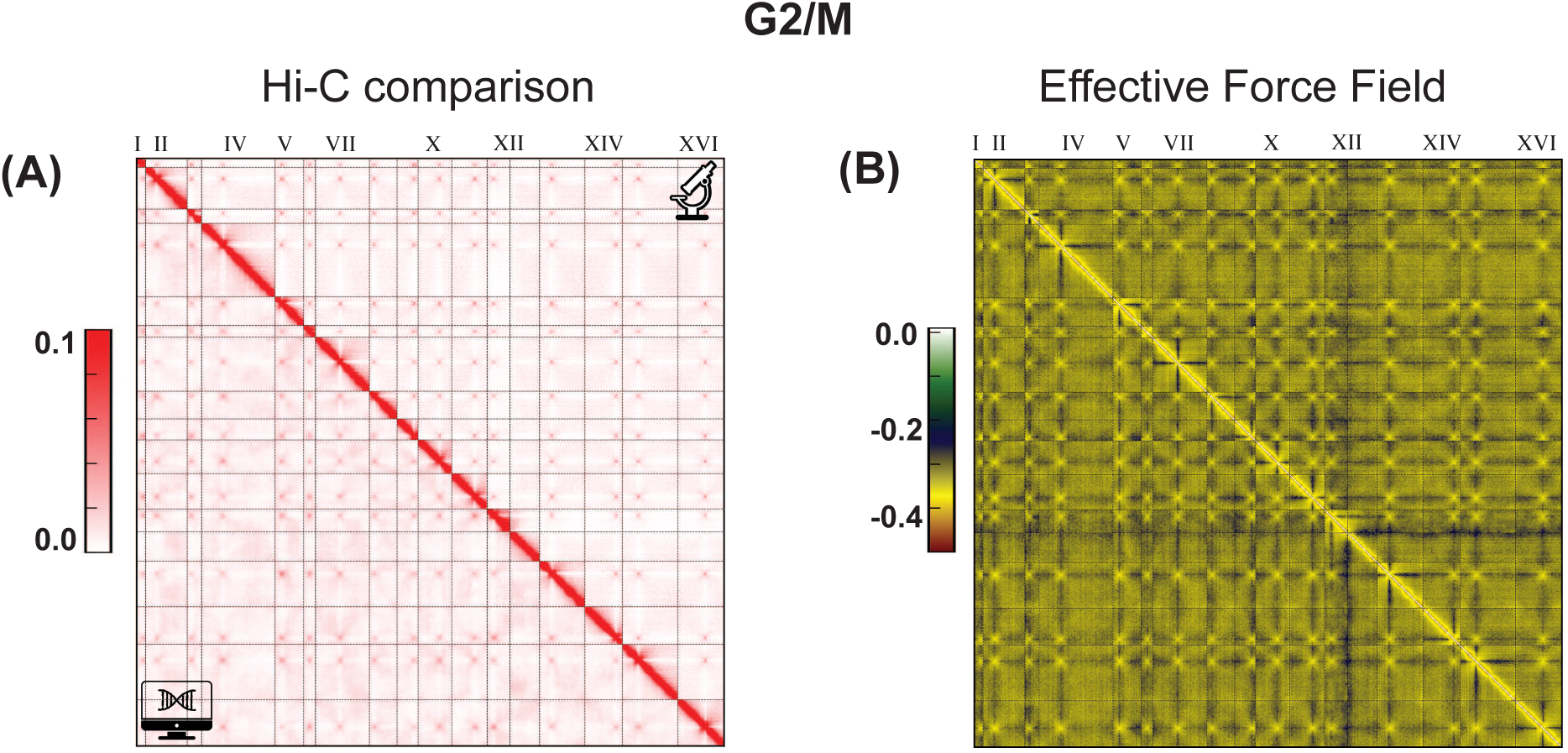
A) Comparison of Hi-C (upper triangle) and simulation (lower triangle) contact maps for G2/M phase. (B) Trained EFF for the G2/M phase obtained using the FI-Chrom method [7] (see Methods for more details). Pairwise energy parameters (*λ*) range from −0.4 (attractive) to 0.4 (repulsive).

**Figure S 4:**
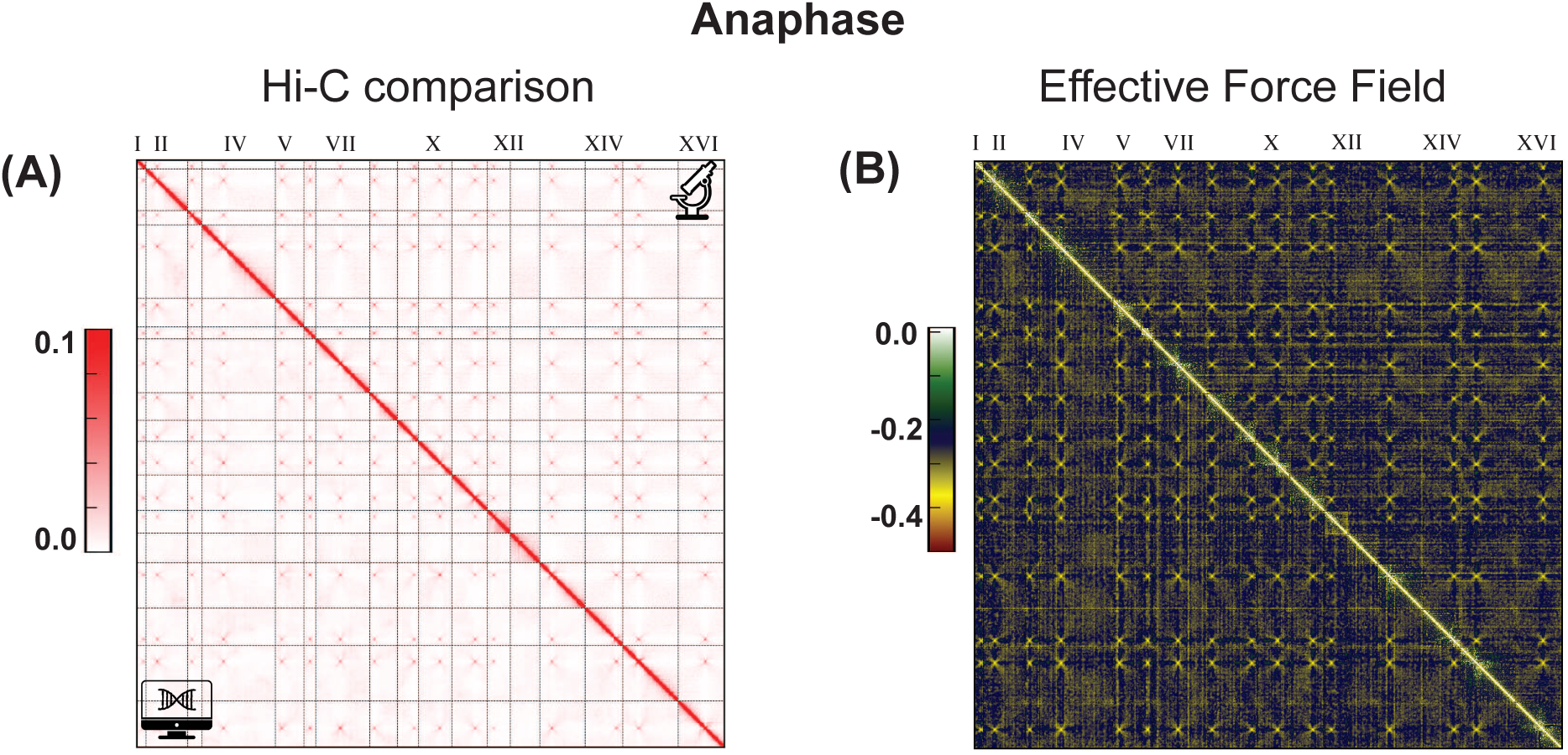
A) Comparison of Hi-C (upper triangle) and simulation (lower triangle) contact maps for Anaphase phase. (B) Trained EFF for the Anaphase phase obtained using the FI-Chrom method [7] (see Methods for more details). Pairwise energy parameters (*λ*) range from −0.4 (attractive) to 0.4 (repulsive).

**Figure S 5:**
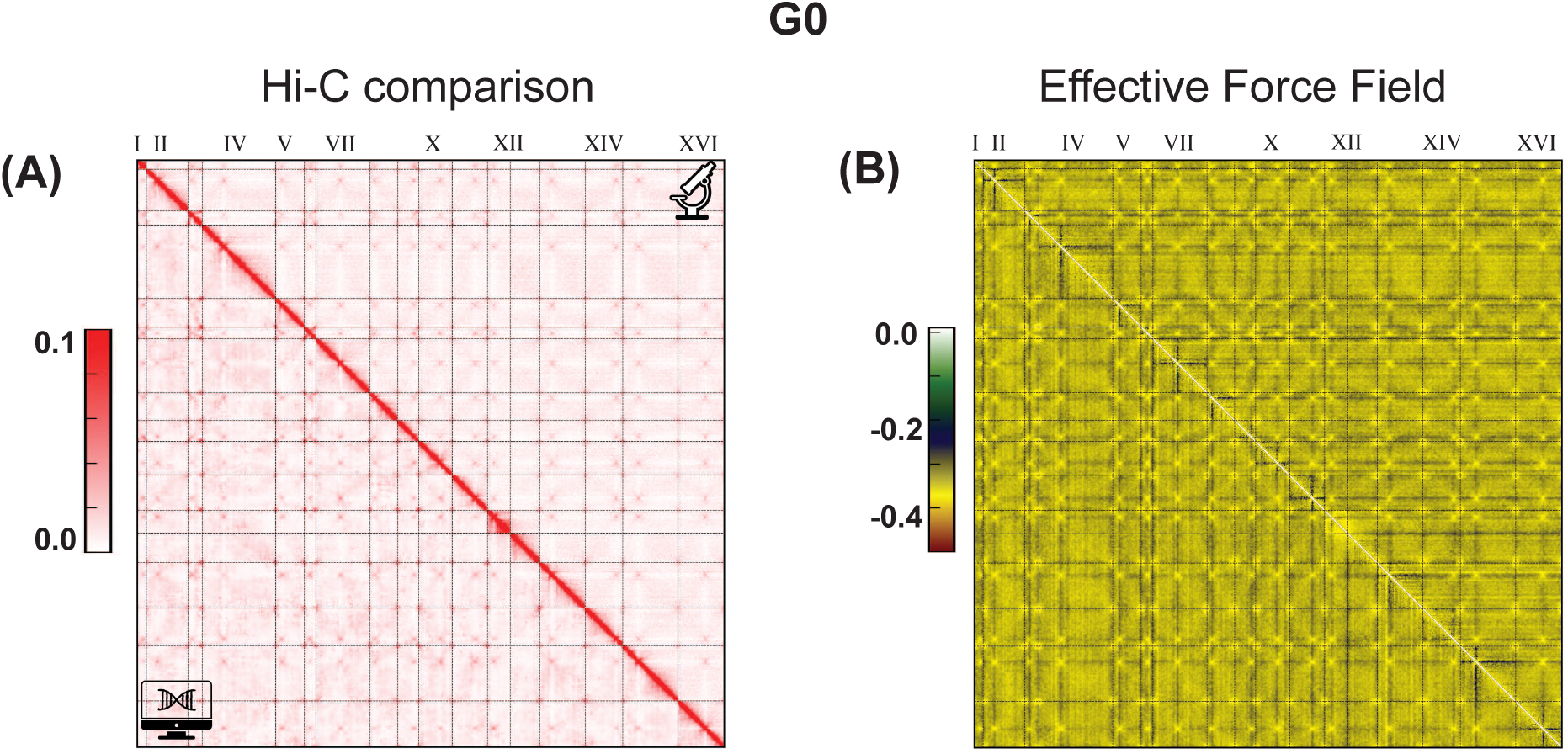
A) Comparison of Hi-C (upper triangle) and simulation (lower triangle) contact maps for G0 phase. (B) Trained EFF for the G0 phase obtained using the FI-Chrom method [7] (see Methods for more details). Pairwise energy parameters (*λ*) range from −0.4 (attractive) to 0.4 (repulsive).

**Figure S 6:**
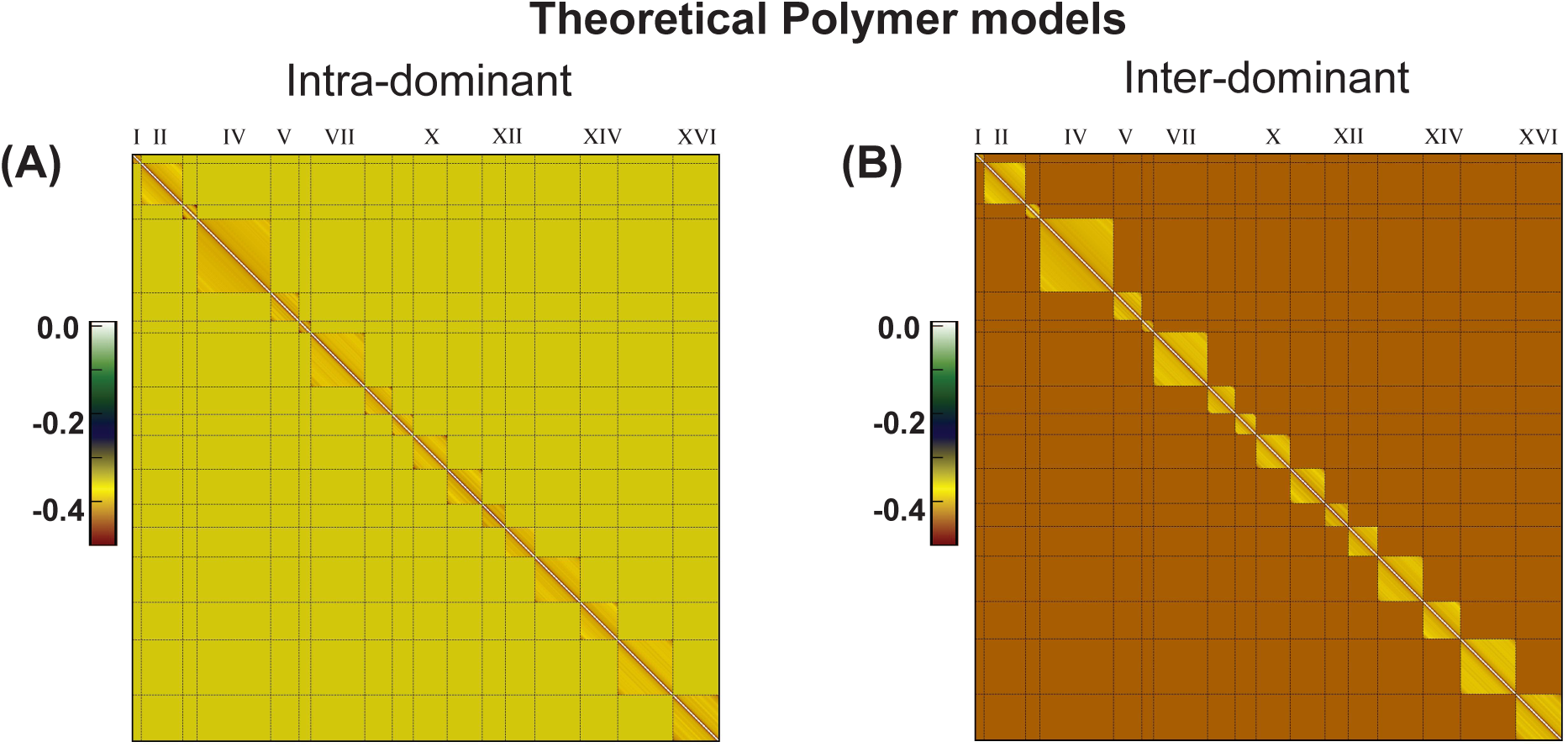
Force fields for the theoretical polymer models. In both models, chromosomes are represented as chains of identical beads. The two models differ only in the relative strength of intra-chromosomal (cis) and inter-chromosomal (trans) bead–bead attractions. (A) Intra-Dominant model, in which the inter-chromosomal (trans) interactions are 20% weaker than the average cis interaction. (B) Inter-Dominant model, in which the trans interactions are 20% stronger than the average cis interaction.

**Figure S 7:**
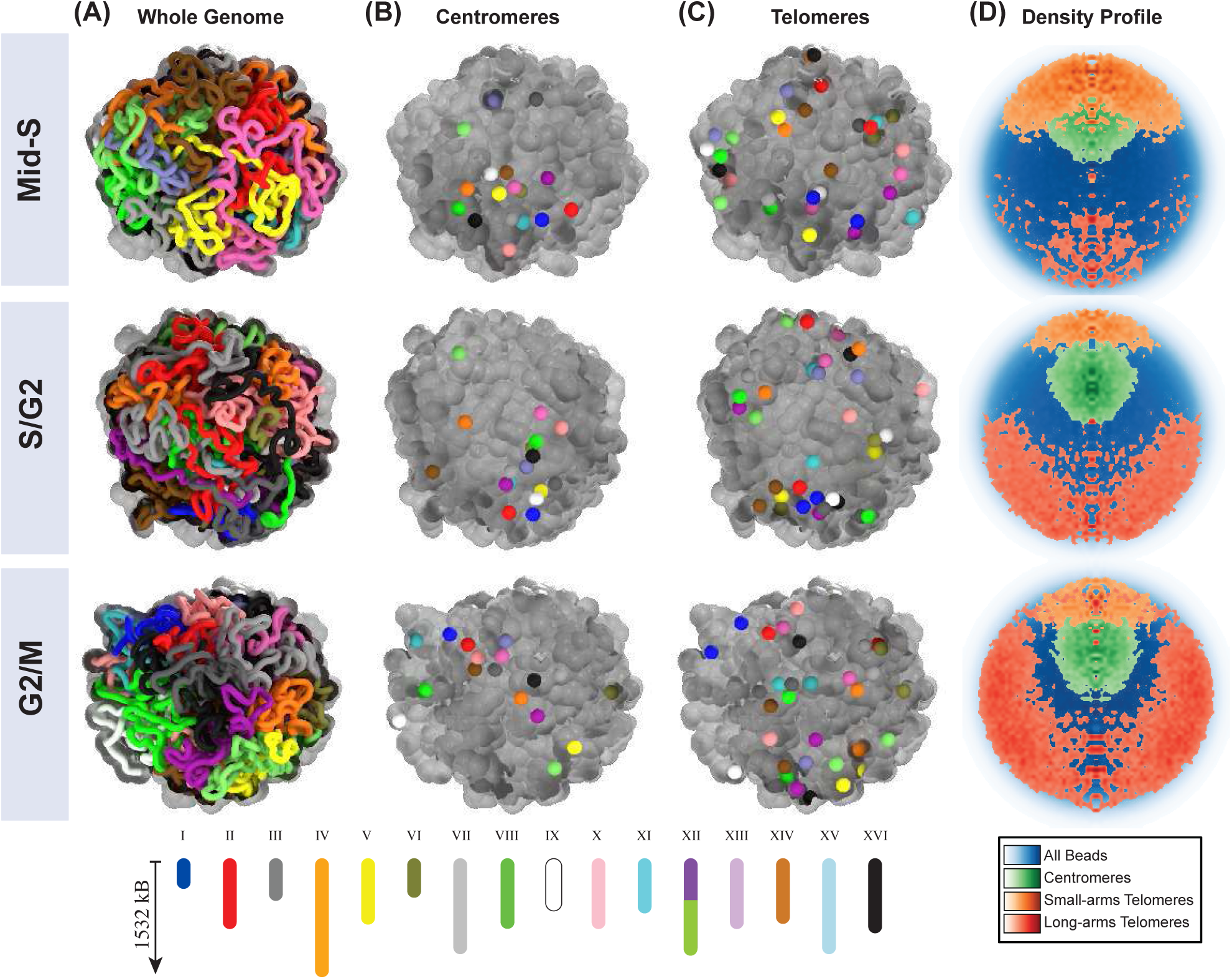
Emergent structural features from the FI-Chrom modeling. Localization profile learned from FI-Chrom modeling for three stages: Mid-S, S/G2, and G2/M. (A) Representative structure of the whole genome, followed by centromere (B) and telomere (C) localization. (D) Density profiles for centromeres, short telomeres, and long telomeres.

**Figure S 8:**
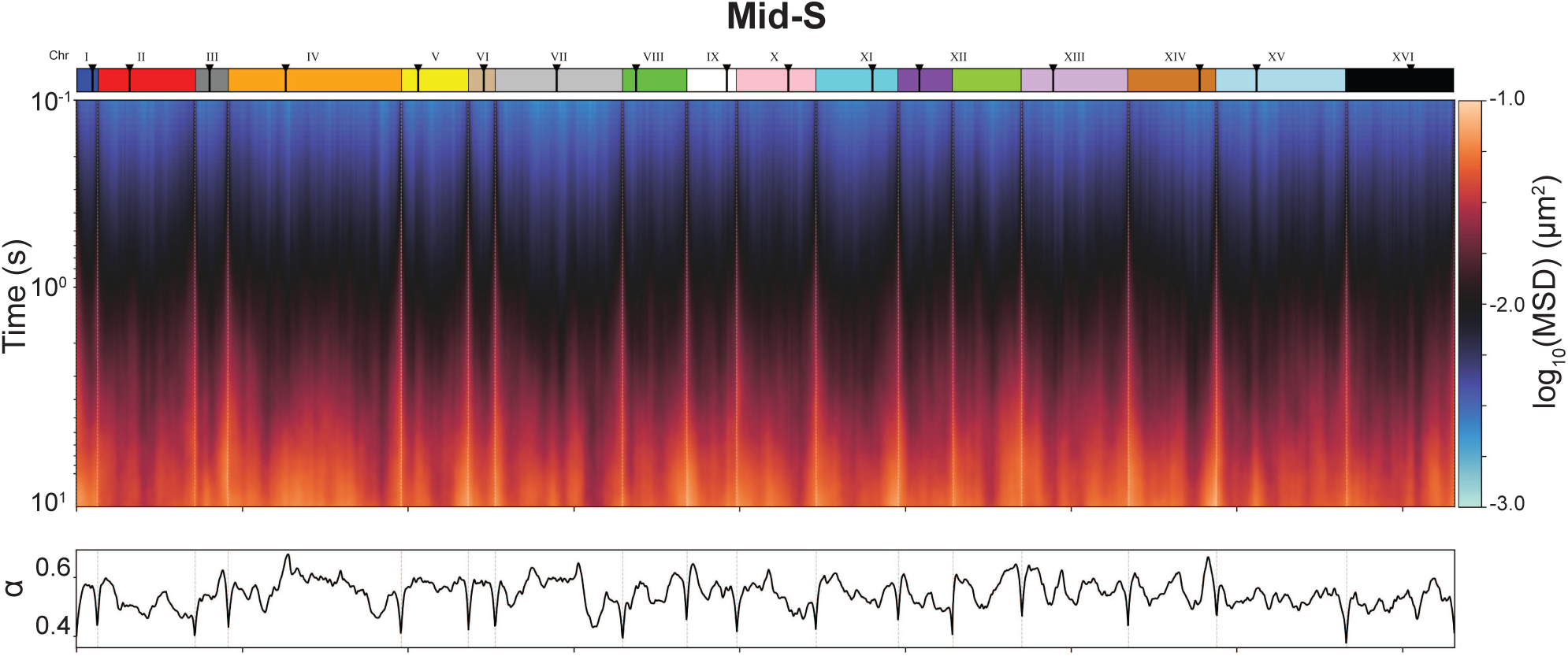
Analysis of the chromatin dynamics. (A) Heatmap showing the *log*_10_(MSD) for each bead (5 kb) in the Mid-S stage. The y-axis represents the lag time in seconds. The centromere regions are marked in the top bar for each chromosome. Below, the MSD scaling exponent (*α*) is calculated for each bead (5 kb).

**Figure S 9:**
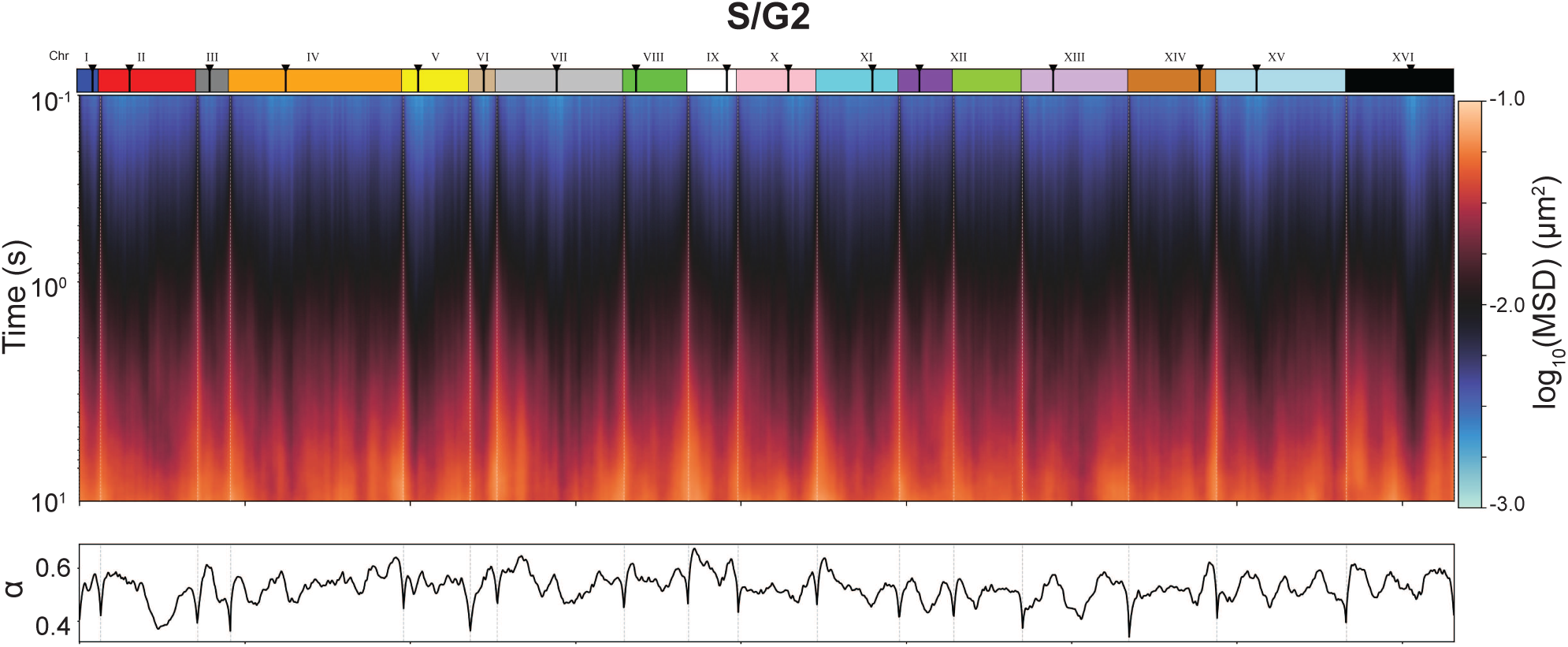
Analysis of the chromatin dynamics. (A) Heatmap showing the *log*_10_(MSD) for each bead (5 kb) in the S/G2 stage. The y-axis represents the lag time in seconds. The centromere regions are marked in the top bar for each chromosome. Below, the MSD scaling exponent (*α*) is calculated for each bead (5 kb).

**Figure S 10:**
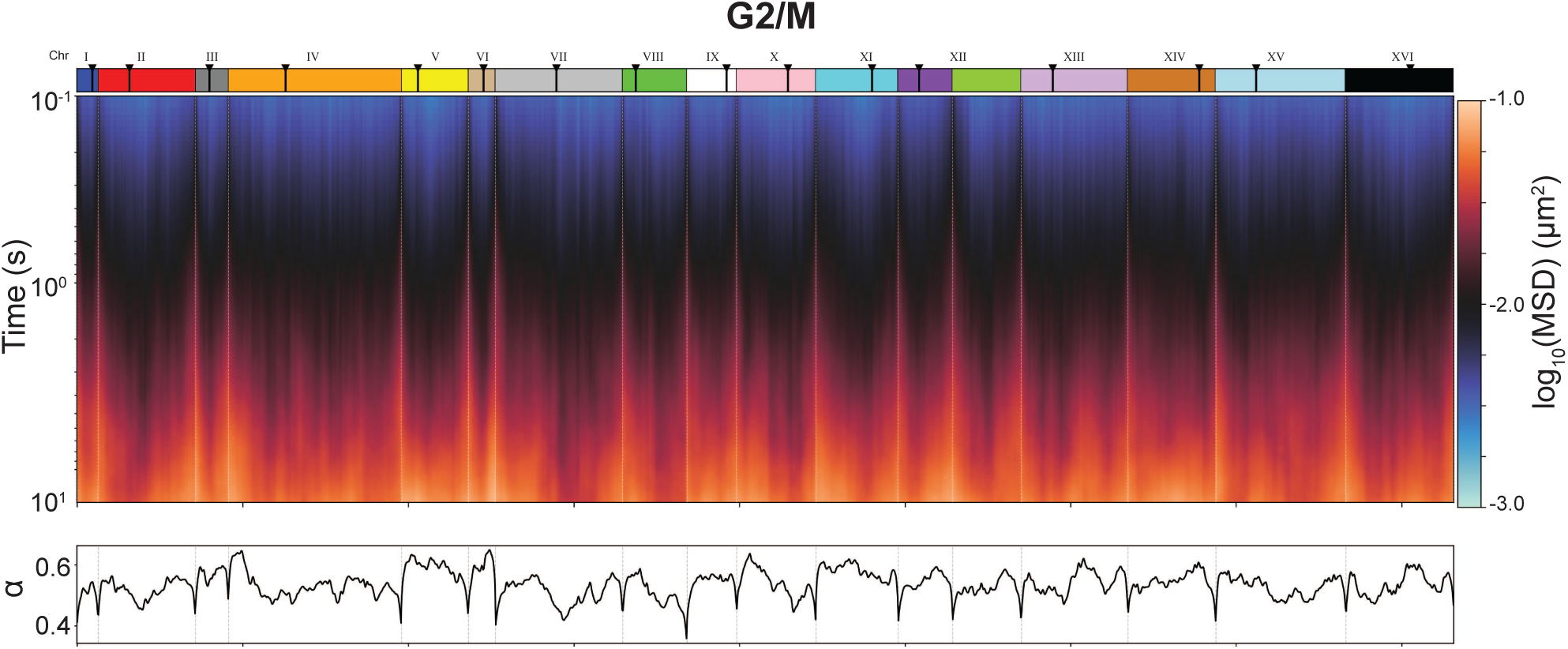
Analysis of the chromatin dynamics. (A) Heatmap showing the *log*_10_(MSD) for each bead (5 kb) in the G2/M stage. The y-axis represents the lag time in seconds. The centromere regions are marked in the top bar for each chromosome. Below, the MSD scaling exponent (*α*) is calculated for each bead (5 kb).

**Figure S 11:**
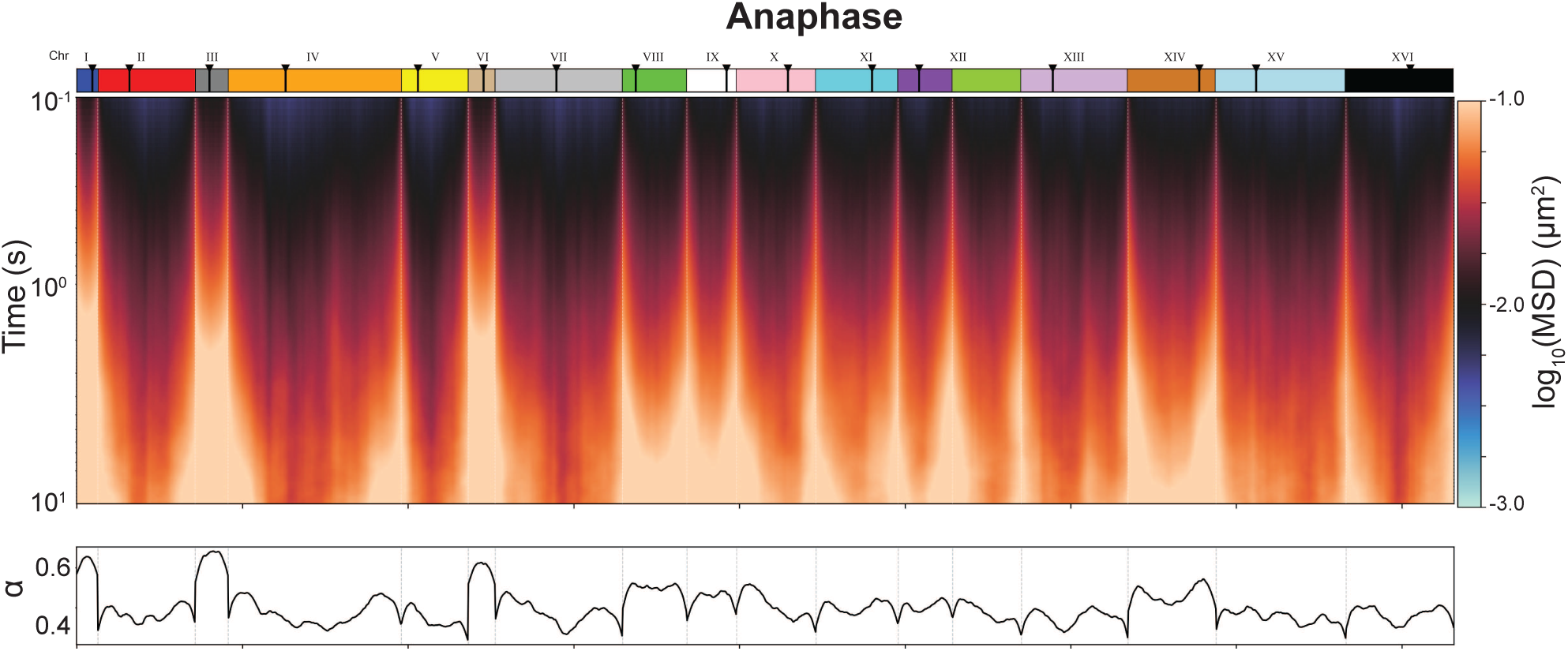
Analysis of the chromatin dynamics. (A) Heatmap showing the *log*_10_(MSD) for each bead (5 kb) in the anaphase stage. The y-axis represents the lag time in seconds. The centromere regions are marked in the top bar for each chromosome. Below, the MSD scaling exponent (*α*) is calculated for each bead (5 kb).

**Figure S 12:**
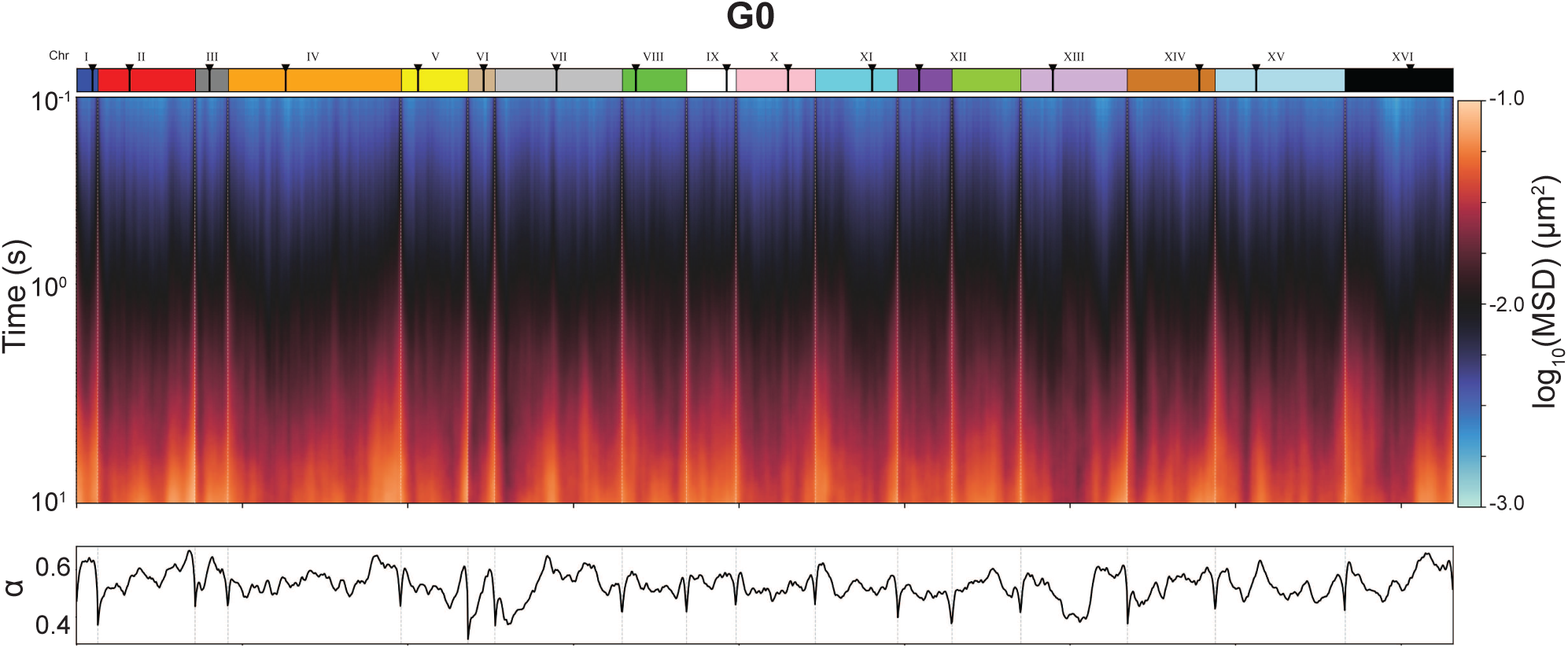
Analysis of the chromatin dynamics. (A) Heatmap showing the *log*_10_(MSD) for each bead (5 kb) in the G0 stage. The y-axis represents the lag time in seconds. The centromere regions are marked in the top bar for each chromosome. Below, the MSD scaling exponent (*α*) is calculated for each bead (5 kb).

